# Ectopic Expression of Distinct *PLC* Genes Identifies ‘Compactness’ as Novel Architectural Shoot Strategy to Cope with Drought Stress

**DOI:** 10.1101/2023.06.02.543233

**Authors:** van Max Hooren, Ringo van Wijk, Irina I. Vaseva, Dominique Van Der Straeten, Michel Haring, Teun Munnik

## Abstract

Phospholipase C (PLC) has been implicated in several stress responses, including drought. Overexpression (OE) of *PLC* has been shown to improve drought tolerance in various plant species. *Arabidopsis* contains nine *PLC* genes, subdivided into four clades. Earlier, OE of *PLC3, -5* or *-7* were found to increase Arabidopsis’ drought tolerance. Here, we confirm this for three other PLCs: *PLC2,* the only constitutively expressed *AtPLC*; *PLC4,* reported to have reduced salt tolerance; and *PLC9,* of which the encoded enzyme was presumed to be catalytically inactive. To compare each *PLC* and to discover any other potential phenotype, two independent OE lines of six At*PLC genes*, representing all four clades, were simultaneously monitored with the GROWSCREEN FLUORO phenotyping platform, under both control- and mild drought conditions. To investigate which tissues were most relevant to achieve drought survival, we additionally expressed At*PLC5* using 13 different cell- or tissue-specific promoters. While no significant differences in plant size, biomass or photosynthesis were found between *PLC* lines and wild-type (WT) plants, all *PLC-OE* lines, as well as those tissue-specific lines that promoted drought survival, exhibited a stronger decrease in convex hull perimeter (= increase in compactness) under water deprivation compared to WT. Increased compactness has not been associated with drought or decreased water loss before, though a hyponastic decrease in compactness in response to increased temperatures has been associated with water loss. We pose that increased compactness leads to decreased water loss and potentially provides a new breeding trait to select for drought tolerance.

## INTRODUCTION

Drought is one of the most impactful stresses in agriculture, affecting 75% of all harvested areas (Boretti & Rosa 2019). Climate change will only make this worse, with increasing periods of high temperature and no rain, as well as a decline in fresh water resources (Kim *et al.,* 2019). Understanding how plants sense and respond to drought is therefore of great importance so that crops can be bred to become more tolerant.

To deal with water deprivation, plants use various strategies (Martignago *et al.,* 2020). The escape strategy induces early flowering, so it can reproduce before dying. Some plants employ the more direct drought avoidance strategy by closing their stomata and by increasing the production of compatible solutes to maintain their water potential, but also by slowing down their life cycle. Lastly, with the drought tolerance strategy, plants are able to improve survival by stimulating DNA repair, ROS scavenging, and expression of Late Embryogenesis Abundant (LEA) proteins and dehydrins that have a chaperone function (Singh *et al*., 2015). The ultimate reaction of plants to water stress is a combination of these strategies, and this may vary between species and ecotypes (Meyre *et al.,* 2001; Bouchabke *et al.,* 2008; Bouzid *et al.,* 2019), and can change in combination with other abiotic- or biotic stresses (Lamaoui *et al.,* 2018; Monohon *et al.,* 2021; Rivero *et al.,* 2021; Sinha *et al.,* 2021)

The initial response to drought may significantly vary, and the reason for this is largely unknown. An interesting candidate in the early sensory mechanism is represented by the lipid signalling pathway headed by phospholipase C (PLC) (Munnik & Vermeer 2010; Munnik 2014; Hong *et al*., 2016; Sagar & Singh, 2021; Ali *et al*., 2022; Fang *et al*., 2023). Overexpression (OE) of *PLC* has been shown to improve drought tolerance in various plant species, including monocots and dicots i.e. maize, rice, tobacco, canola, and Arabidopsis (Wang *et al.,* 2008; Georges *et al.,* 2009; Tripathy *et al.,* 2012; Zhang *et al.,* 2014; van Wijk *et al.,* 2018; Zhang *et al.,* 2018a; Zhang *et al.,* 2018b; Deng *et al.,* 2019; Chen *et al.,* 2021).

In animal systems, PLC is a well-established signalling enzyme that becomes activated in response to receptor stimulation by various signals (Katan & Cockcroft, 2020). Upon activation, PLC hydrolyses the minor plasma membrane phospholipid, phosphatidylinositol 4,5- bisphosphate (PIP_2_) into two second messengers: inositol 1,4,5-trisphosphate (IP_3_) and diacylglycerol (DAG). While the water-soluble IP_3_ diffuses into the cytosol where it triggers the release of Ca^2+^ from the ER by specific binding of a IP_3_ gated-Ca^2+^ channel, DAG remains at the plasma membrane where it recruits and activates members of the protein kinase C (PKC) family. The subsequent increase in cytosolic Ca^2+^ concentration and change in phosphorylation status of various target proteins, trigger a host of downstream responses, affecting numerous cellular processes (Katan & Cockcroft, 2020).

The PLC-signalling system has been found to be strongly conserved in flowering plants, though the main downstream targets, i.e. the IP_3_ channel and PKC, are typically absent (Munnik & Testerink 2009; Munnik 2014). Instead, flowering plants seem to use the phosphorylated products of the PLC hydrolysis as second messengers, i.e. inositol polyphosphates (IPPs; including IP_5_, IP_6_, IP_7_ and IP_8_) and phosphatidic acid (PA), which are generated by IPP kinases (IPKs) and DAG kinases (DGKs), respectively (Munnik *et al.,* 2000; Munnik & Vermeer 2010; Gillaspy 2013). Plant targets for IP_5_ or IP_6_ include ligand-gated Ca^2+^channels, proteins involving mRNA transport (e.g. GLE1) and F-box proteins involved in auxin- and jasmonate perception (*e.g.* TIR1 and COI1) (Munnik & Testerink 2009; Heilmann 2016a; Noack & Jaillais 2017). Similarly, IP_7_ and IP_8_ have been implicated in phosphate signalling through binding SPX domains (Williams *et al.,* 2015; Laha *et al.,* 2016; Ried *et al.,* 2021). Targets for PA include protein kinases, phosphatases, small G-proteins, transcription factors, and membrane transporters, involving processes such as endocytosis, membrane trafficking, organisation of the cytoskeleton, osmoregulation, and gene expression (Munnik 2001; Wang *et al.,* 2006; Li *et al.,* 2009; Testerink & Munnik 2011; McLoughlin & Testerink 2013; Pleskot *et al.,* 2013; Pokotylo *et al.,* 2014; Thomas & Staiger 2014; Hou *et al.,* 2016; Ufer *et al.,* 2017; Pokotylo *et al.,* 2018; Yao & Xue 2018; Wang *et al.,* 2019; Kolesnikov *et al.,* 2022).

Animal PLCs predominantly hydrolyse PIP_2_ but can also hydrolyse its immediate precursor, phosphatidylinositol 4- monophosphate (PIP; Levin *et al.,* 2017). The quantities of these minor lipids are approximately equal, 1:1. In flowering plants, however, PIP_2_ levels are ∼30-100 fold lower while PIP concentrations are similar to those in animals (Munnik *et al.,* 1998; Meijer & Munnik 2003; Heilmann 2016b; Noack & Jaillais 2017). So, for plants, it is more likely that PLC hydrolyses PIP in response to stimulation (Munnik, 2014). There are, however, conditions in which plants locally and temporarily increase their PIP_2_ levels. I.e. upon heat- or osmotic stress, during cell division, at the tip of growing root hairs and pollen tubes, and in response to polyamines, which likely reflects PIP_2_’s role as lipid second messenger itself (van Leeuwen *et al.,* 2007; Ischebeck *et al.,* 2010; Simon *et al.,* 2014; Bloch *et al.,* 2016; Zarza *et al.,* 2020). During these circumstances, PLC may have an additional function, i.e. to attenuate PIP_2_ signalling besides initiating IPP- and PA signalling.

How overexpression of *PLC* leads to increased drought tolerance is unknown. The fact that it works in both monocots and dicots indicates that the mechanism is conserved (Wang *et al.,* 2008; Georges *et al.,* 2009; Tripathy *et al.,* 2012; Zhang *et al.,* 2014; van Wijk *et al.,* 2018; Zhang *et al.,* 2018a; Zhang *et al.,* 2018b; Deng *et al.,* 2019; Chen *et al.,* 2021. Van Hooren *et al*., 2023). While metazoan PLCs can be grouped into six subfamilies, which are characterized by conserved domains that explain their regulation by e.g. G-protein coupled receptors or receptor tyrosine kinases, plant PLCs, lack most of such domains and only contain the minimum present in all eukaryotic PLCs, i.e. a catalytic X- and Y-domain that fold over the lipid headgroup, an EF-hand, and C2 domain; the later explaining the Ca^2+^ sensitivity of the enzyme (Munnik 2014). Plant PLCs mostly resembles PLCζ, which is the only isoform that lacks the pleckstrin homology (PH) domain responsible for binding PIP_2_. Metazoan PLCζ is specifically expressed in sperm cells, and it is still unknown how this isoform is regulated (Thanassoulas *et al*., 2022), as is the case for plant PLCs (Munnik 2014; Hong *et al*., 2016; Sagar & Singh, 2021; Ali *et al*., 2022; Fang *et al*., 2023).

The Arabidopsis genome encodes nine *PLC* genes, which can be subdivided into four clades (Hunt *et al.,* 2004; Tasma *et al*., 2008). Earlier, OE of *PLC3*, PLC4; *PLC5* and *PLC7* have each been found to increase the tolerance of Arabidopsis plants to water stress (van Wijk *et al.,* 2018; Zhang *et al.,* 2018a; Zhang *et al.,* 2018b; Van Hooren *et al*., 2023). *PLC2* is the only constitutively expressed *PLC* and is also highest in expression of all *AtPLCs* (Tasma *et al*., 2008). The fourth PLC clade is represented by *PLC8* and *PLC9*, of which the encoded proteins were predicted to be catalytically inactive due to certain mutations in the catalytic X-domain (Hunt *et al.,* 2004), though experimental evidence for this is lacking. Interestingly, *PLC9-OE* were reported to contain higher IP_3_ levels under heat stress while *plc9* knock out (KO) mutants exhibited reduced levels in response to heat (Zheng *et al.,* 2012). Moreover, *AtPLC9* overexpression increased the thermotolerance of both Arabidopsis and rice (Zheng *et al.,* 2012; Liu *et al.,* 2020), while plc9 mutants were more sensitive to heat stress. These experimental results imply that AtPLC9 does have activity.

Promoter-GUS fusions in Arabidopsis showed that *PLC* expression is most prominent in vascular tissue in the shoot (*PLC1, -2, -3, -5* and *-7*), and either in the vasculature of developing root and root primordia (*PLC3,-5*) or throughout the whole root (*PLC1, -2, -7*). Expression has also been found in the root tip (*PLC2, -7*), stomata (*PLC2, -5*), and trichomes (*PLC2, -3, -5 and -7*) (Hunt *et al.,* 2004; Kanehara *et al.,* 2015; van Wijk *et al.,* 2018; Zhang *et al.,* 2018a; Zhang *et al.,* 2018b).

*PLC* overexpression is typically achieved with POLYUBIQUITIN 10 (UBQ10)- or 35S promoters that ectopically and constitutively express genes to high amounts (Grefen *et al.,* 2010). Experiments performed with such lines have mainly focussed on survival under severe drought stress, comparing the tolerance of OE lines of a single *PLC* gene with that of WT (Wang *et al*., 2008; Georges *et al*., 2009; Tripathy *et al*., 2012; Deng et al., 2018; van Wijk *et al*., 2018; Zhang *et al*., 2018a; Zhang *et al*., 2018b). In this study, the GROWSCREEN phenotyping platform (Jansen *et al.,* 2009) was used to quantitatively compare the behaviour of Arabidopsis *PLC-OE* lines from all four *PLC* clades under mild water stress conditions, i.e. *PLC2*, *PLC3*, *PLC4*, *PLC5*, *PLC7* and *PLC9*, in comparison with wild type (WT) plants. To investigate which tissues were most relevant for the *PLC* induced-drought tolerance, we constructed 13 different ’*Tissue-Specific Expression of PLC5’* (*TSEP*) lines, and took these along in the phenotyping screen. The GROWSCREEN-FLUORO setup allowed for daily measurements of plant growth, architecture, colour and chlorophyll fluorescence, measuring responses to water deprivation in a non-lethal fashion (Osmolovskaya *et al.,* 2018). Both early and late responses to water deprivation were monitored, as well as the recovery after the post-stress irrigation of the plants. Instead of just estimating the final outcomes of the imposed water deprivation, an overview of the progress of the stress may provide new information on the plant’s response, following the dynamics of the studied parameters.

## RESULTS

### PLC2-OE promotes drought stress tolerance

Of all Arabidopsis *PLC* genes, *PLC2* is expressed the highest and is the only *PLC* that is constitutively expressed and not induced by stress (Hunt *et al.,* 2004; Tasma *et al.,* 2008). To test whether OE of this *PLC* would also promote drought tolerance, distinct *PLC2*-*OE* lines were generated using *pUBQ10* and *p2x35S* promoters. Enhanced expression was validated using qPCR (∼5-17 fold) (Fig. 1A) and Western blot analysis of the two lines with the highest gene expression (Fig. 1B). A clear increase in a ∼66 kD protein, the expected size of PLC2, was observed as wells as two minor bands at ∼37 and ∼29 kD, which add up as 66 kD, likely representing PLC2- breakdown products. The same *PLC-OE* lines revealed an increase in drought tolerance after 18 days of withholding water (Fig. 1C).

**Figure 1.**
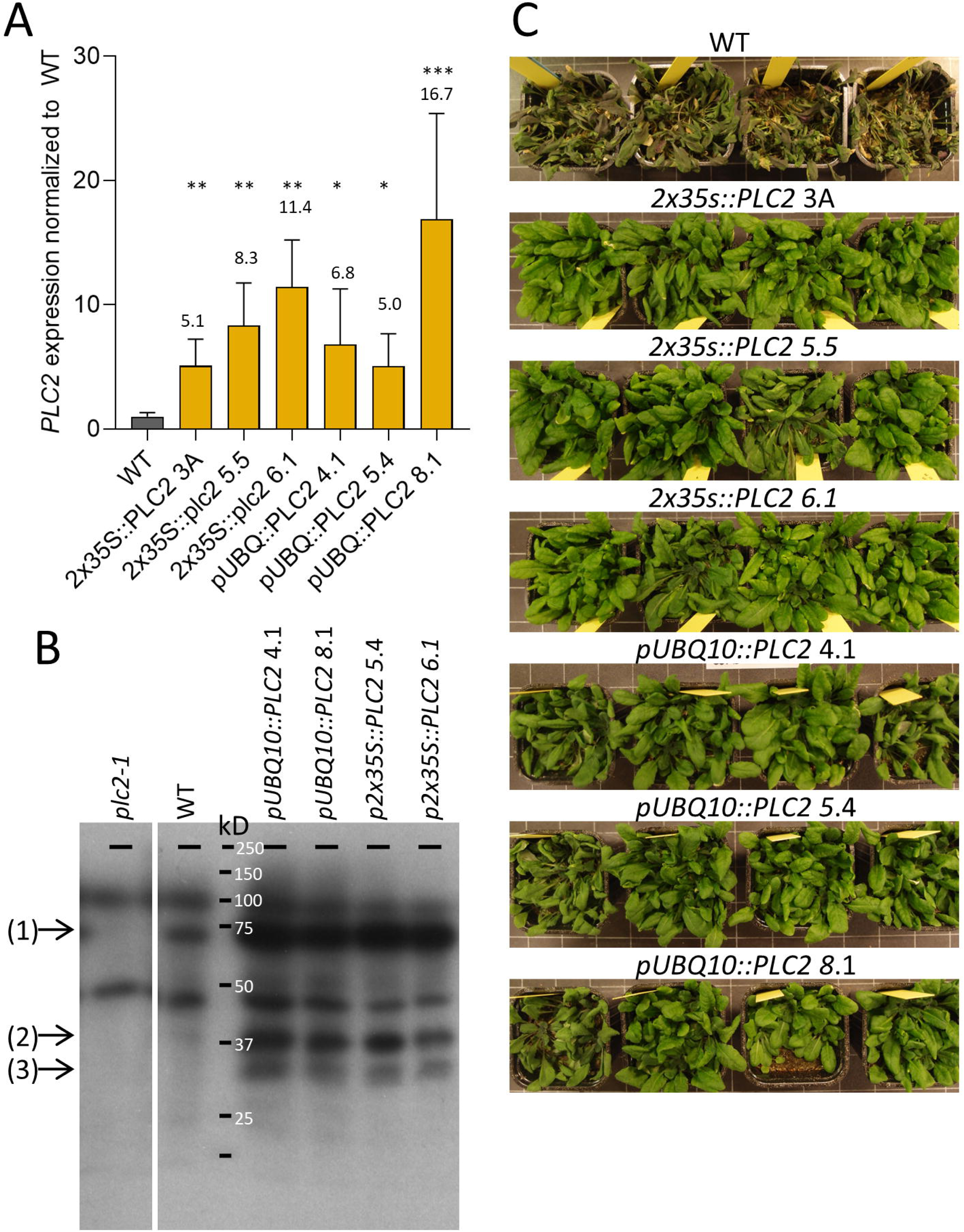
survival rates of *PLC2*-OE lines under water stress. A) *PLC2* expression in WT, OE lines as measured by qPCR. Values are normalized to *SAND* and WT and represent the means with SD of three-four biological replicates. (ANOVA test, *= P<0.05; **= P<0.01; ***= P<0.001). B) PLC2 detection using Western blot with anti-AtPLC2 1/2000 (D’ambrosio 2017) and GARPO 1/5000 on 11-day old roots of WT, KO and OE lines of PLC2. (1) AtPLC2:3xmVenus ∼147 kD, (2) AtPLC2 ∼66 kD, (3) Breakdown product AtPLC2 ∼37 kD, (4) Breakdown product AtPLC2 ∼29 kD C) Survey of plant survival of WT and *PLC2-*OE lines after water stress in a single experiment. 37-day-old plants were water deprived for 18 days, after which they were watered again, and photographs were taken after 9 days.

### Overexpression of presumed-inactive PLC9 increases drought tolerance too

In the literature, PLC8 and PLC9 are suggested to be catalytically inactive (Hunt *et al.,* 2004; Tasma *et al.,* 2008), though experimental evidence is lacking for this. To investigate this in more detail, the catalytic domains of PLC8 and PLC9 were compared *in silico* with those of the other Arabidopsis PLCs as well as with rat PLCδ1 (*Rattus norvegicus*), of which the crystal structure has been resolved (Essen *et al.,* 1997) and several essential amino acids for activity have been identified (Ellis *et al.,* 1998). Except for PLC8 and PLC9, all Arabidopsis PLCs contained the same key amino acids in the catalytic site as rat PLCδ1 (Supplemental Table S3). Mutations in the catalytic side for PLC8 and PLC9 were H311L, H356P, E390K, K440R and Y155R, of which the two histidine residues are located at prime positions within the catalytic centre, as revealed by modelling of the catalytic sites of PLC5 and PLC9, and using the protein structure of rat PLCδ1 as template (Fig. 2A). Mutations of these two histidines in rat PLCδ1, resulted in a dramatic reduction of PIP_2_ binding, i.e. by 6000-fold for His^311^ and 20.000-fold for His^356^ (Ellis *et al.,* 1998). While these results may indicate that PLC8 and PLC9 are theoretically inactive, PLC8 and PLC9 also gained two new cationic amino acids (i.e. E390K, Y155R) that may take over. Especially, since PIP is the more likely substrate of plant PLCs than PIP_2_ (Munnik, 2014)

**Figure 2.**
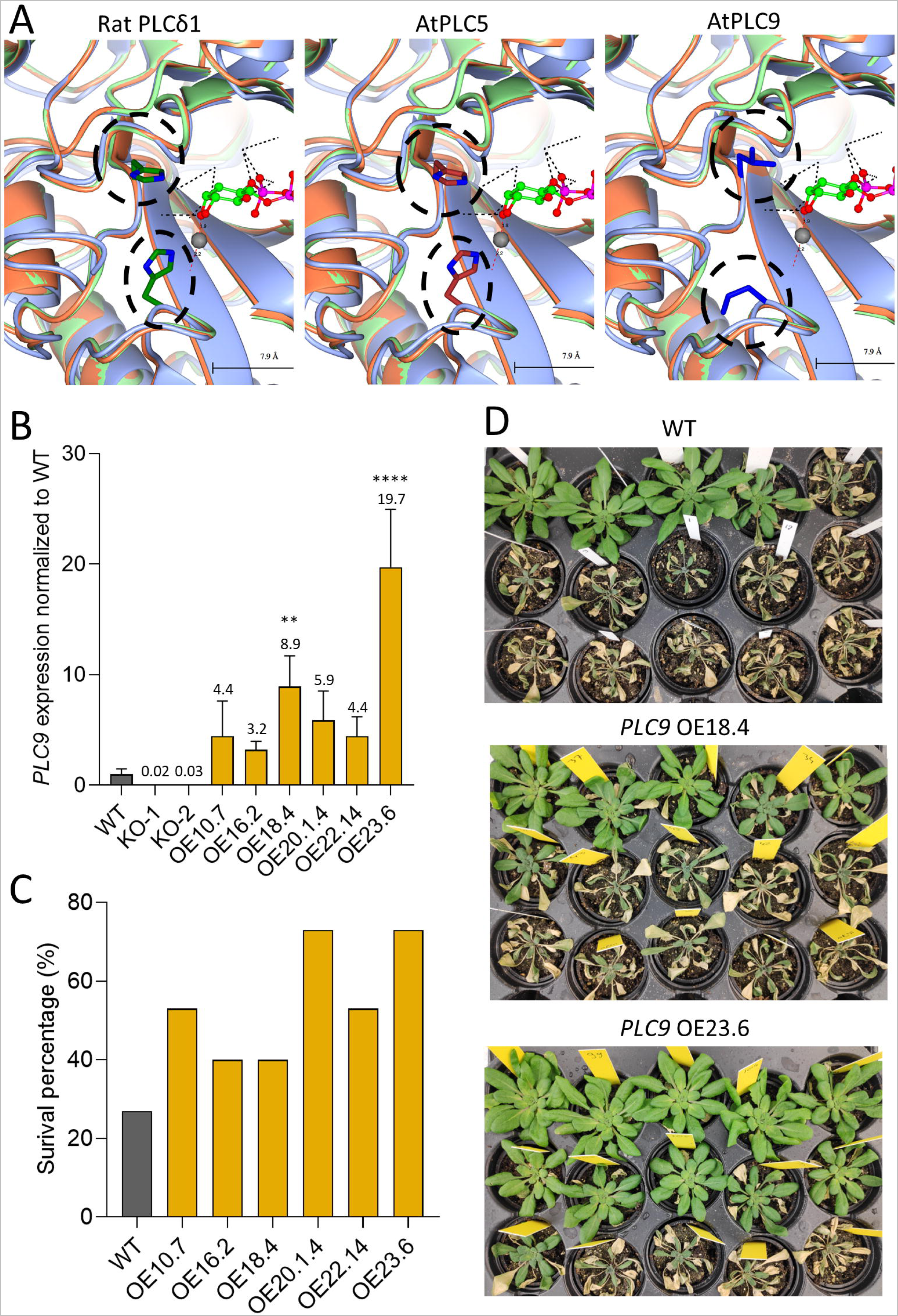
Structure modelling of catalytic site of PLC9, and survival rates of *PLC9*-OE lines under water stress. A) Protein structures of Arabidopsis PLC9 compared to AtPLC5 and Rat PLCδ (*1DJX*). Encircled are the His residues missing from PLC9. B) *PLC9* expression in WT, KO- and OE lines as measured by qPCR. Values are normalized to *SAND* and represent the means with SD of three-four biological replicates. (ANOVA test, **= P<0.01; ****= P<0.0001). C) Survey of plant survival of WT and *PLC9-*OE lines after water stress in a single experiment. Three weeks-old plants were water deprived for 3 weeks, after which they were watered again D) Photographs at the plants used in figure 1C after three days of rewatering (pots with surviving plants were grouped together per line).

A number of independent *PLC9*-OE lines were generated, and their *PLC9* expression monitored in homozygous T3 plants using qPCR, along with two *plc9*-KO lines to validate the specificity of the primers (*PLC8* is a very close homolog). As shown in Figure 2B, all *PLC9-OE* lines revealed an increase in *PLC9* expression (∼3- 20 fold), and these lines also showed increased survival rates after three weeks of drought stress (Figs. 2C, D).

### Shoot architecture of PLC-OE under water deprivation

For the subsequent large GROWSCREEN phenotyping experiment, *PLC*-OE lines with the highest overexpression, were selected (i.e. pUBQ10 lines #4.1 and #8.1 for *PLC2* and #18.4 and #23.6 for *PLC9*), together with two independent OE lines of *PLC3, -4, -5,* and *-7*.

Plant rosettes were monitored during control- and water deprivation conditions between 25-38 days after sowing (DAS) and during recovery, until harvesting at 45 DAS. Since all *PLC*-OE lines exhibited increased survival during water deprivation, we were specifically interested in traits that would distinguish *PLC-OE*s from WT plants. Using a PCA plot, traits were visualized that differentiated *PLC-OE*s from WT under control (Fig. 3A) and drought conditions (Fig. 3B). In order to correct for age, and thus size of the plants, data from both PCA plots was normalized per day, for each day at control conditions, or during the period of water deprivation for the stressed plants. Arrow length and -direction indicate how much and to which principal component (PC) each trait contributed. Data from the measuring devices, i.e. RGB and fluorescence, were separated into distinct PCA plots to facilitate their analysis (Supplemental Fig. S1). RGB measurements were taken at more time points, and PCA results showed greater explanatory power than the fluorescence measurements. Hence, we focussed on the results obtained from RGB measurements. Data from the fluorescence measurements can be found in Supplemental Figures S2-12.

**Figure 3.**
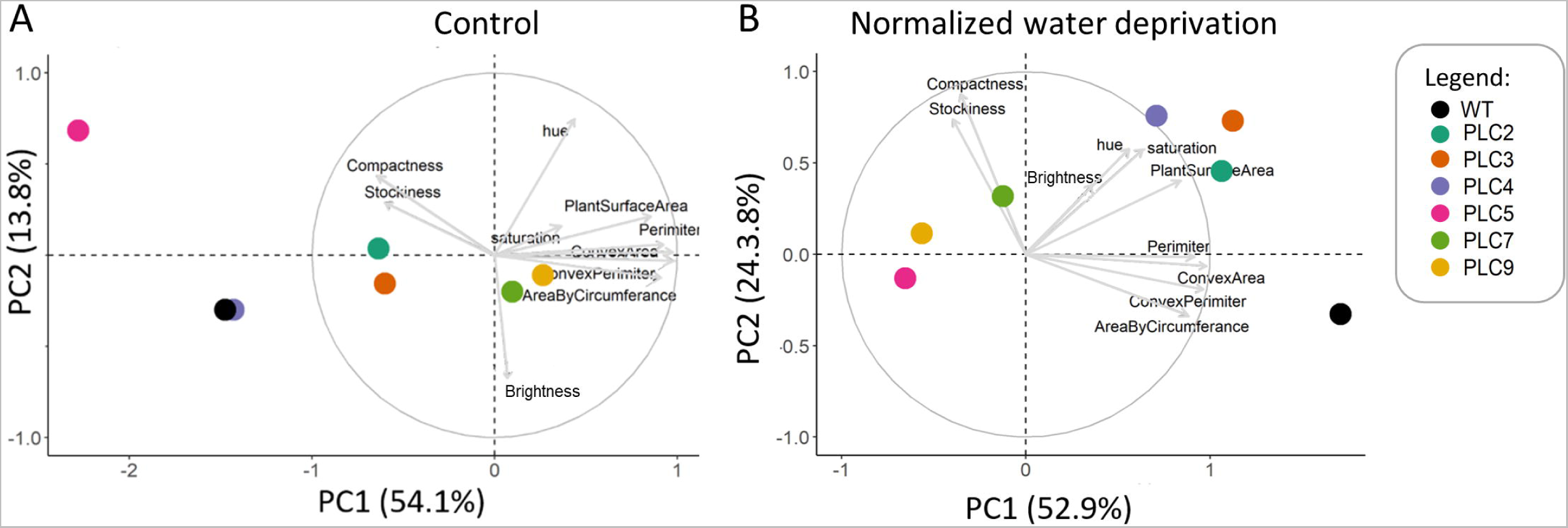
PCA plots showing altered plant morphology responses of *PLC-OE* lines compared to WT in their response to water stress. PCA plots from control- (left panel, 21-45 DAS) and water-limiting conditions normalized to control (right panel; 30-38 DAS (day 5-13 of stress). Data was normalized per day and per trait, to account for plants being larger at later days. Arrows size and direction indicate how much each trait contributes to each principal component (PC). Percentages of total variance represented by PC1 and PC2 are shown in parentheses. N = 40-60

At control conditions, no specific traits were found that would distinguish *PLC-OE*s from WT (Fig. 3A). *PLC-OE* lines acted differently from each other and from WT, which is in line with earlier observations that OE of *PLC* did not result in striking growth phenotypes (van Wijk *et al.,* 2018; Zhang *et al.,* 2018a). Only *PLC5*-OE plants were found to be smaller and more compact, confirming earlier observations (Zhang *et al.,* 2018b).

When looking at the effect of water deprivation, *PLC-OE* lines did group together and, more importantly, distinguished themselves from WT (Fig. 3B). The traits in this PCA plot can roughly be divided into three groups that clustered together: I [*Perimeter*, *Convex hull perimeter*, *Area by circumference* and *Convex hull area*], II [*Compactness and Stockiness*], and, III [*Hue*, *Saturation*, *Brightness value* and *Projected Plant surface area*] (Fig. 3B).

*Projected plant surface area* represents the total number of pixels in RGB that was identified as plant (Fig. 4A). Under control conditions, OE of *PLC3, -7* and *-9* exhibited a statistically significant larger *Projected plant surface area* than WT. As mentioned earlier, *PLC5-OE* lines were slightly smaller than the WT, but at control conditions this was not statistically significant. Under water deprivation, the changes between WT and *PLC-OE* lines became smaller and were no longer significantly different (Fig. 4B), except for *PLC5*-OE that was significantly smaller. By normalizing the *Projected plant surface area* of water deprived data to the control conditions, we saw that *PLC5-, PLC7-* and *PLC9*-OE lines exhibited a significantly stronger growth decrease than WT (Fig. 4C).

**Figure 4.**
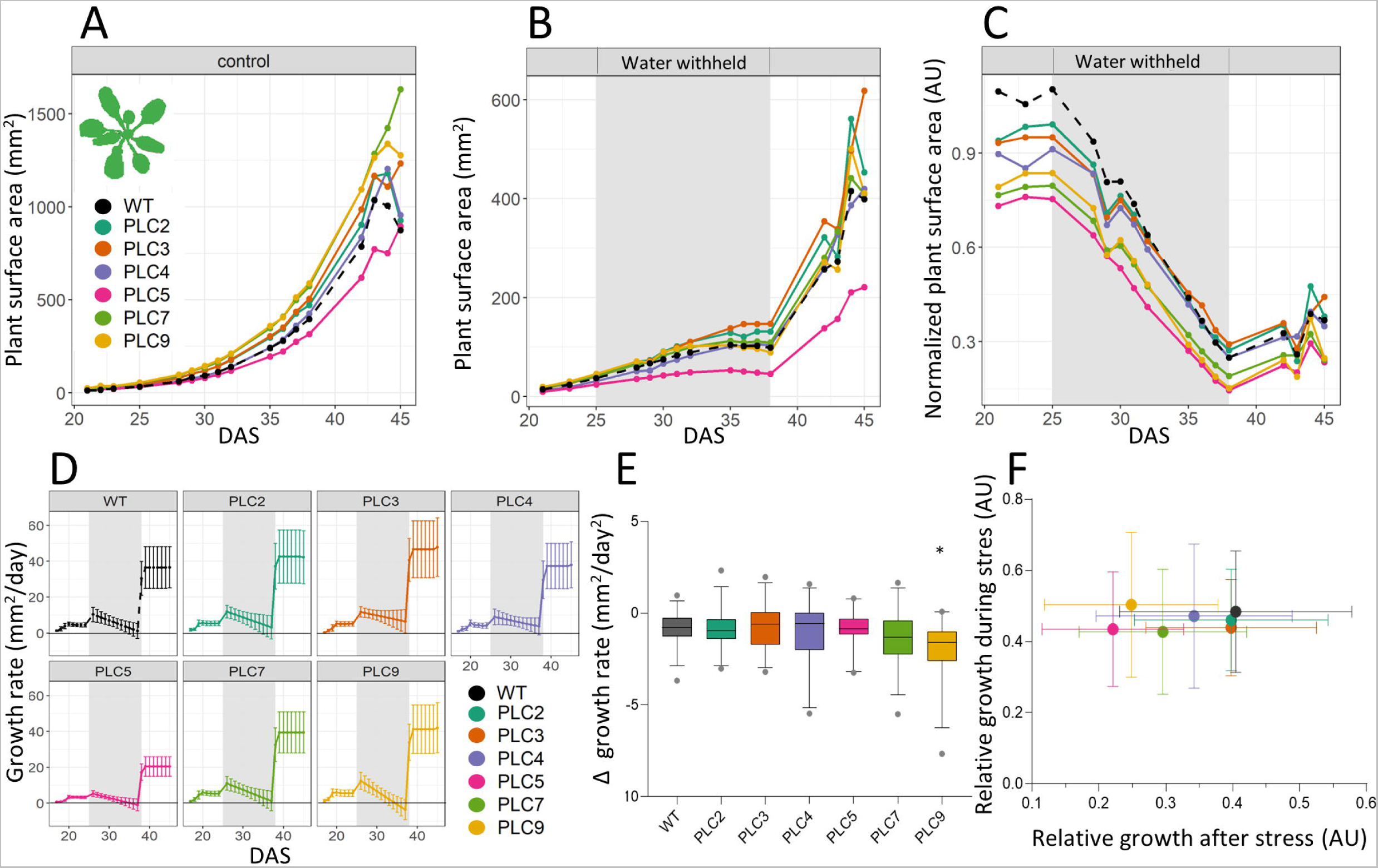
*PLC* overexpression does not change Projected plant surface area under water limiting conditions. Arabidopsis WT and *PLC-OE* lines plants were monitored using the GROWSCREEN-FLUORO system for indicated times. For each genotype, two different insertion lines with 30 replicates were used. A) Projected leaf area (schematically drawn in upper left corner) at control conditions. B) Projected leaf area at water-withholding conditions (25-37 DAS). C) Projected leaf area at water-withholding conditions normalized to control. D) Modelled growth rates at water-deprivation conditions. Each individual plant was modelled, before, during and after drought stress, and then used to determine the relative growth rate per day. E) Growth rate declines during water deprivation. Modelled growth rates for each plant during the water withholding period were made using the formula: ax^2^+bx+c formula, where a was extracted as an indication of how fast growth declined in response to drought. Error bars represent a 95% confidence interval. Data was analysed by one-way ANOVA followed by a Dunnet’s test. Statistically significant differences with WT are indicated by * (p<0,05) F) Ratio of growth during and after water withholding, normalized to growth during and after withholding at control conditions. Error bars represent SEM.

The growth rate was determined to investigate how well the different lines continued to grow during water stress. However, as sowing dates were spread across three days, and measuring was performed at the same time, not all plants could be compared at the same DAS. To overcome this, growth curves were plotted for each individual plant. This modelled data could then be used to determine growth rate per day (Fig. 4D). As the growth per day is very much correlated to the initial size of the plant, we also determined the derivative of the ’growth per day’, or ’decline in growth per day’ for the time during water withholding (Fig 4E). Of the different lines, only *PLC9*-OE plants exhibited a statistically significant bigger growth-rate decline. Hence, we conclude that the drought-tolerant phenotype of *PLC-OE*s is not caused by a difference in growth response. Growth relative to control, during and after water-withholding, was plotted to see whether a different growth strategy could be observed for the *PLC-OE*s (Fig. 4F). Growth during water deprivation mirrored the data from Figure 4C, but during recovery, no differences between WT and *PLC-OE*s were observed.

The parameters in the PCA plots, that allowed differentiation between WT and *PLC-OE*s were: *Convex hull perimeter*, *Plant perimeter*, *Convex hull area* and *Area by circumference*. These traits seemed to have a similar influence in separating the plant lines, and also contributed greatly to the difference between WT and *PLC-OE*s. The *Convex hull perimeter* (Fig. 5) is determined by drawing a line across the extremities of each leaf to get the perimeter of the plant, which is different from the *Plant perimeter* (Supplemental Fig. S13) that follows the exact contours of the leaves. One might expect the *Plant perimeter* to be more precise, but values highly fluctuate as leaves tend to overlap with each other as plants grow older. C*onvex hull area* (Fig. 6) uses the lines drawn by the *Convex hull perimeter* to calculate the area. The *Area by circumference* (Supplemental Fig. S14) takes the longest leaf and draws a circle using that leaf length as the radius for the circle. In both cases, the convex hull data were more consistent, with less outliers, and revealing the same general patterns. Hence, we focused on the convex hull data. Both *convex hull perimeter* and *-area* were slightly larger in *PLC-OE*s under control conditions, with the exception of *PLC5-*OE. For OE lines of *PLC3, -7* and -*9* this was statistically significant (Figs. 5A, 6A). However, this difference disappeared during water deprivation, with *PLC5*-OE becoming significantly smaller than WT (Figs. 5B, 6B). That the *PLC-OE*s lose their slightly larger *Convex hull perimeter* & *- area*, and thus respond more strongly to the water deprivation, is clearly visible when normalizing the water deprived data to control (Figs. 5D, 6D), and was statistically significant for *PLC5-, 7-* and *9*-OE lines.

**Figure 5.**
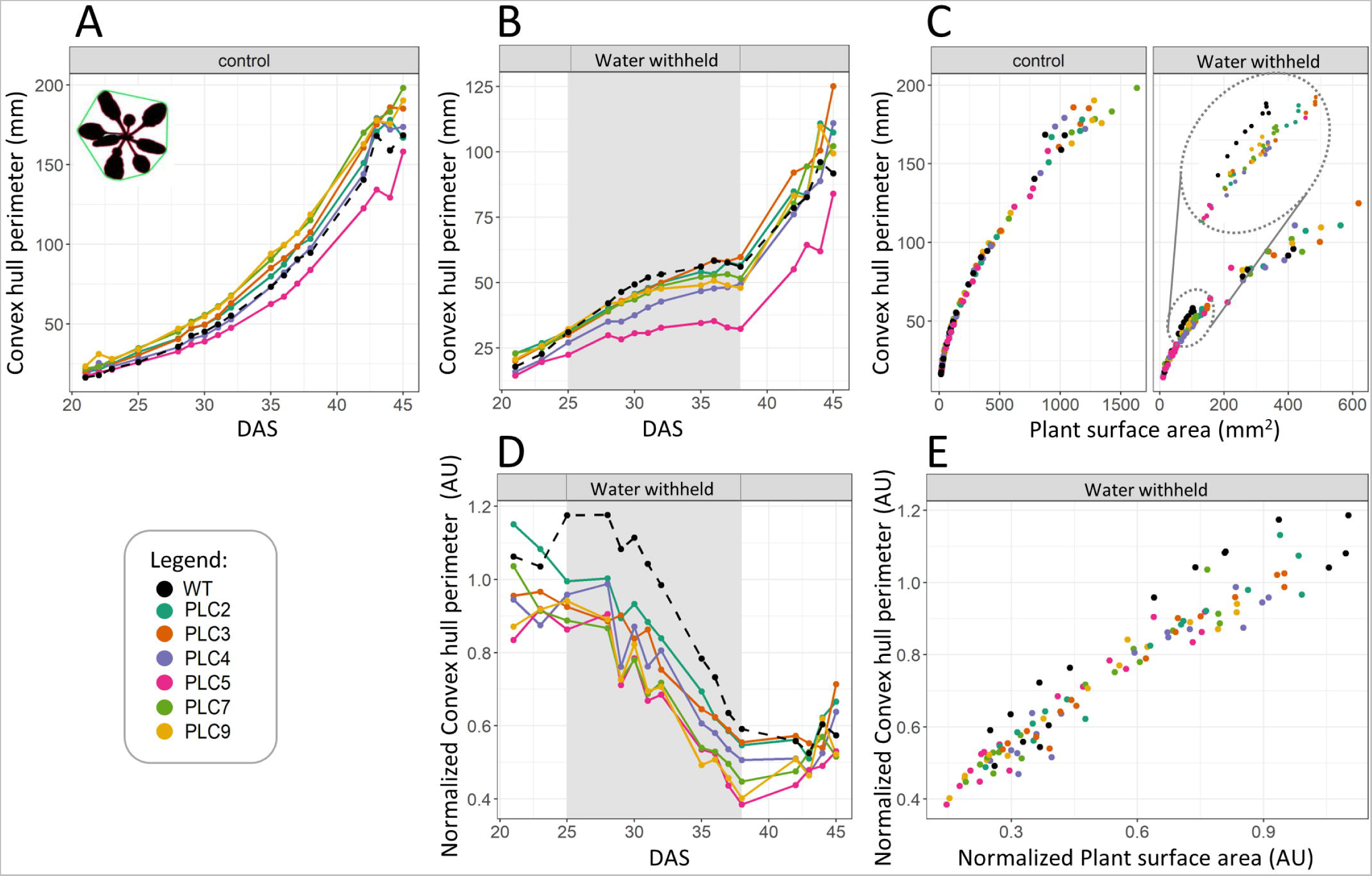
PLC overexpression leads to a decrease in convex hull perimeter in response to water stress. The convex hull perimeter (schematically indicated in the upper left corner of panel A) was measured in: A) Plants grown at control conditions. B) Plants at water-withholding conditions, with grey shading indicating period of water withholding (25-38 DAS). C) Water-withholding conditions normalized to control per day D) Water withholding conditions normalized to control conditions. E) Water withholding conditions normalized to Projected plant surface area.

**Figure 6.**
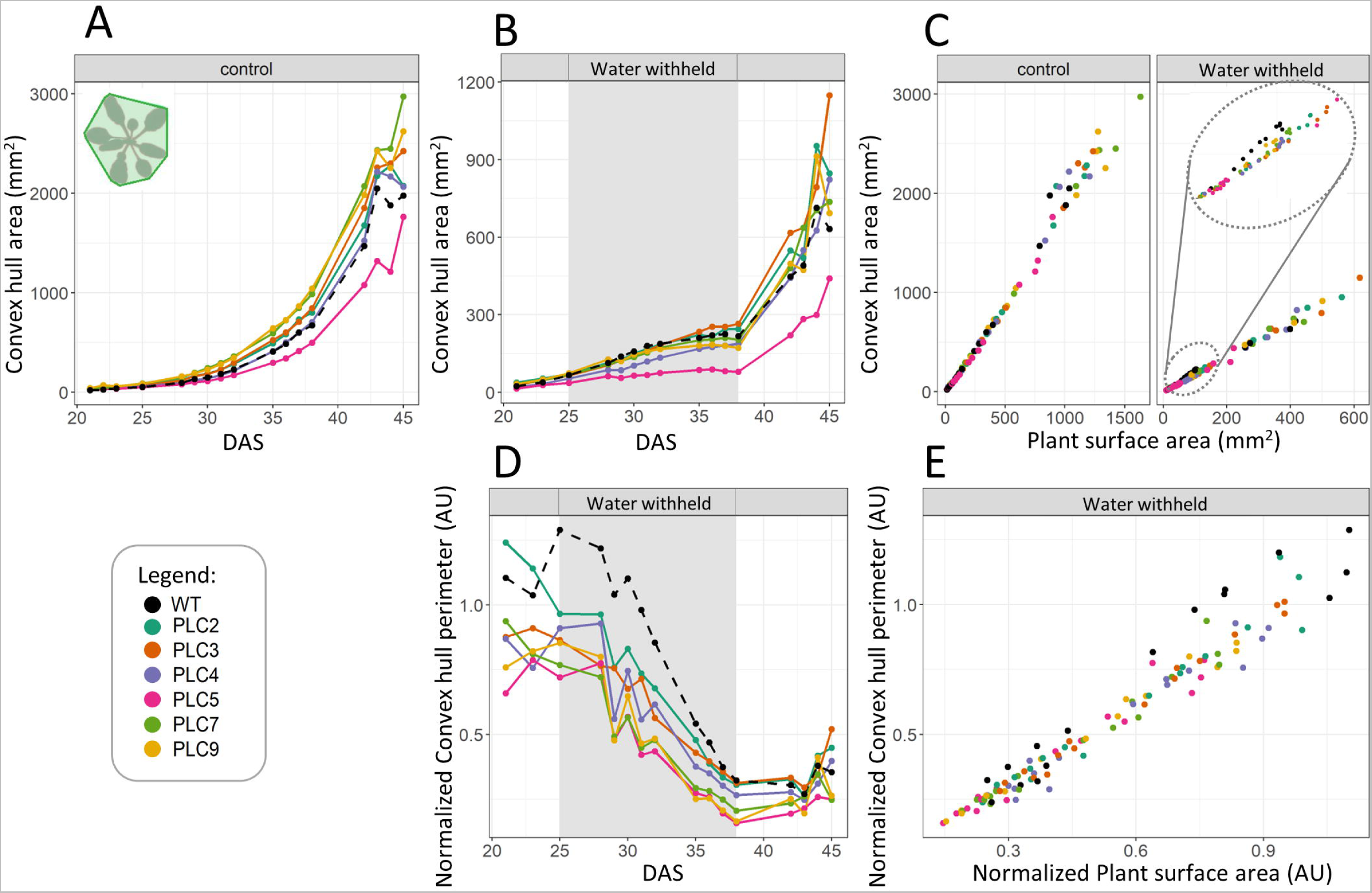
*PLC-OE* plants exhibit decreased convex hull area in response to water stress. A) Convex hull area (indicated in upper left corner panel A) at control conditions. B) Convex hull area at water withholding conditions (25-38 DAS). C) Projected plant surface area plotted against convex-hull area. D) Convex hull area water withholding conditions normalized to control conditions. E) The normalized water withholding data Projected plant surface area data plotted against the convex hull perimeter data.

When comparing the *Convex hull perimeter* & -*area* to *Projected plant surface area* (Figs. 5C, 6C) under control conditions, no substantial differences were observed. However, under water-limited conditions, when comparing plants of equal size, the *PLC-OE* lines tend to have smaller *Convex hull perimeter*. The same, although to a lesser extent, was found for the *Convex hull area*. This was most clearly observed in the *Projected plant surface area* range of 50-100 mm^2^, which corresponds to the average size of WT plants during water deprivation. When normalized to control conditions, it became clear that *PLC-OE*s had a greatly reduced *Convex hull perimeter* & -*area* compared to WT, with the change being statistically significant for *PLC5, -7* and -*9* OEs throughout the whole stress period, while the rest of the PLCs exhibited only significant values at intermittent points (Supplemental Table S4).

The relation between *Projected plant size* and *Convex hull area* can be expressed by *Compactness* (Fig. 7). *Compactness* is the *Projected plant surface* area divided by the *Convex hull area*, so it gives the ratio of how much of the *Convex hull area* is actually covered by the rosette. Similarly, plant size can be divided by *Area by circumference*, resulting in the *Stockiness* of a plant. However, as *Area by circumference* had a higher variation than the *Convex hull area*, *Stockiness* also had a higher variation than *Compactness* (Supplemental Fig. S15).

**Figure 7.**
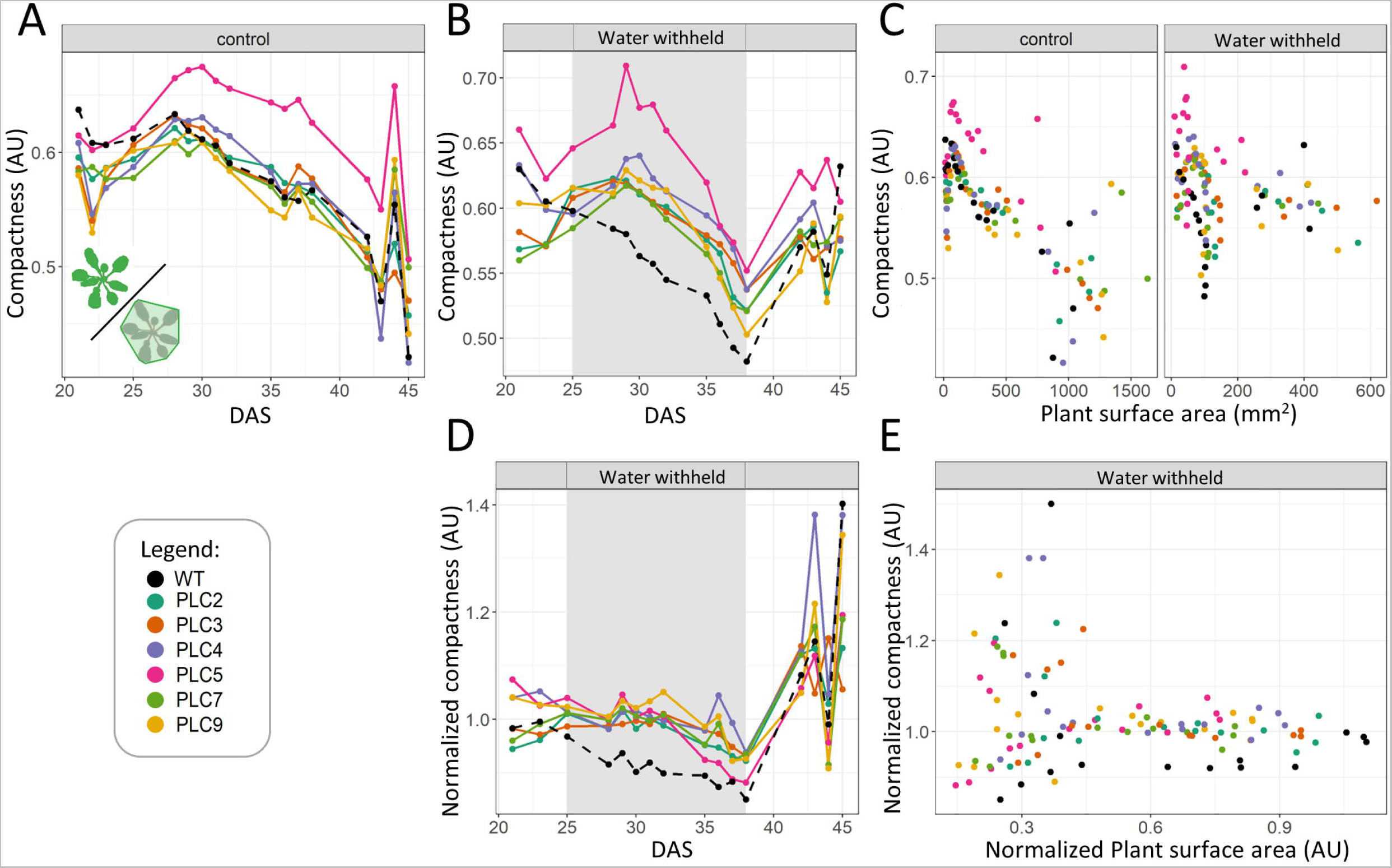
*PLC-OE* lines have increased compactness in response to water limitation Compactness is schematically indicated in green in the upper left corner of panel A. A) Compactness at control conditions. B) Compactness at water withholding conditions, with grey shading indicating the particular period (25-38 DAS). C) Compactness under control and water-withholding conditions. For each measurement day, Projected plant surface area is plotted against compactness. D) Compactness at water withholding conditions normalized to control conditions. E) The normalized water withholding data Projected plant surface area data plotted against the compactness data.

At control conditions, *Compactness* reached a maximum at 28-30 DAS in all plants. Only *PLC5*-OE plants were significantly more compact than WT at control conditions (Fig. 7A). During water limitation, WT plants became less compact as stress prolonged. Strikingly, *PLC-OE*s retained their *Compactness* as at control conditions during the most part of the drought treatment, and only decreased their compactness at the very last days of stress (Figs. 7B, 7D). As a group, the *PLC-OE*s were statistically more compact under, and in their response to, water stress than WT. Comparing *Plant size area* to *Compactness* mainly showed that no matter the size of the plant, *PLC5*-OE was always more compact (Fig. 7C), and that *PLC-OE*s remain more compact relative to their size in response to drought (Fig. 7E).

### Chlorophyll content and fluorescence of PLC-OE plants under water deprivation

In this large-scale experiment, chlorophyll content was not directly measured, but indirectly using the *Hue* of a plant, which is correlated with its chlorophyll content (Majer *et al.,* 2010; Liang *et al.,* 2017; Farago *et al.,* 2018). Since a calibration curve was lacking, only the relative chlorophyll content could be measured.

*Hue*, *Saturation* and *Brightness* were determined by transforming RGB values of the area covered by the *Projected plant surface area*. Under control conditions, *Hue* steadily increased over time, levelling off at the latest time points, indicating the greening of plants as they grow older (Supplemental Fig. S16A). Water-deprived plants initially still increased their *Hue*, but this rapidly decreased after 6-8 days of water withholding, and increased again after rewatering (Supplemental Fig. S16B). Only *PLC5*-OE plants showed a significant difference with WT at control conditions, being slightly greener at the later timepoints. Aside from this observation, no significant differences between *PLC-OE*s and WT were observed in either condition, nor after normalization, or when compared to *Projected plant surface area* (Supplemental Figs. S16C, S16D, S16E). Similarly, *Brightness* (Supplemental Fig. S17) and *Saturation* (Supplemental Fig. S18) revealed no difference between WT and *PLC-OE* lines. Interestingly, *Saturation* responded very quickly to water deprivation, as 4 days after the start of the stress period already a decline was observed, while at control conditions the *Saturation* remained constantly increasing.

Photosynthetic activity is often measured using the *F_v_/F_m_*ratio, which indicates how much of the photoreactive centres present are actually available for photosynthesis (Yao *et al.,* 2018). Under stress conditions, this ratio is typically reduced to limit the production of ROS (Dalal & Tripathy 2018). Under control conditions, we observed that the *F_v_/F_m_* ratios remained more or less stable (Supplemental Fig. S6A), but indeed went down after seven days of water stress (Supplemental Fig. S6B). In both conditions, however, no statistically significant difference between WT and *PLC-OE* lines was found. After normalization, some significant changes were found during the first couple of days of water deprivation, but this faded as stress prolonged (Supplemental Fig. S6C).

### Biomass of PLC-OE plants after water deprivation

After 13 days of water deprivation, normal watering was resumed again. After a week of recovery, plants were harvested and *Fresh weight* (FW) (Supplemental Figs. S19A, S19B) and *Dry weight* (DW) (Supplemental Figs. S19D, S19E) of the shoots determined. The *Relative Water Content* was determined using the formula: (FW-DW)/FW*100%) (Supplemental Figs. S19G, S19H). At control conditions, plants overexpressing *PLC3-, -7* and *-9* were significantly heavier than WT, while *PLC5*-OE plants were significantly lighter than WT, in both FW- and DW. The *Relative water content* at control conditions was only significantly different for *PLC5*-OE. After water stress, only *PLC9-* and *PLC3*-OE lines were significantly different in FW- and DW compared to WT, and no differences in the *relative water content* among the *PLC-OE* lines was recorded. Normalizing the water-stressed plants to their control counterparts revealed that none of the lines responded significantly different from WT to water stress (Supplemental Figs. S19C. S19F, S19I), which is in strong contrast to their survival rate in response to drought.

### Phenotyping Tissue specific-PLC5 (TSEP) lines

To investigate which cells/tissues are most important in gaining drought survival through *PLC* expression, a set of cell- and tissue-specific promoters was used to drive expression of *PLC5* (Supplemental Table 2). This was based on earlier individual experiments in which *PLC5-OE* appeared to generate the strongest drought tolerance (not shown). Normally, *PLC5* is expressed in the vasculature (phloem), hydathodes and guard cells (Zhang *et al*., 2018b). Thirteen so-called *Tissue-Specific Expression of PLC5* (*TSEP*) lines were generated. An overview of the tissue specificity of the used promoters and a schematic representation of the expression of each line is given in Figures 8A & 8B.

**Figure 8.**
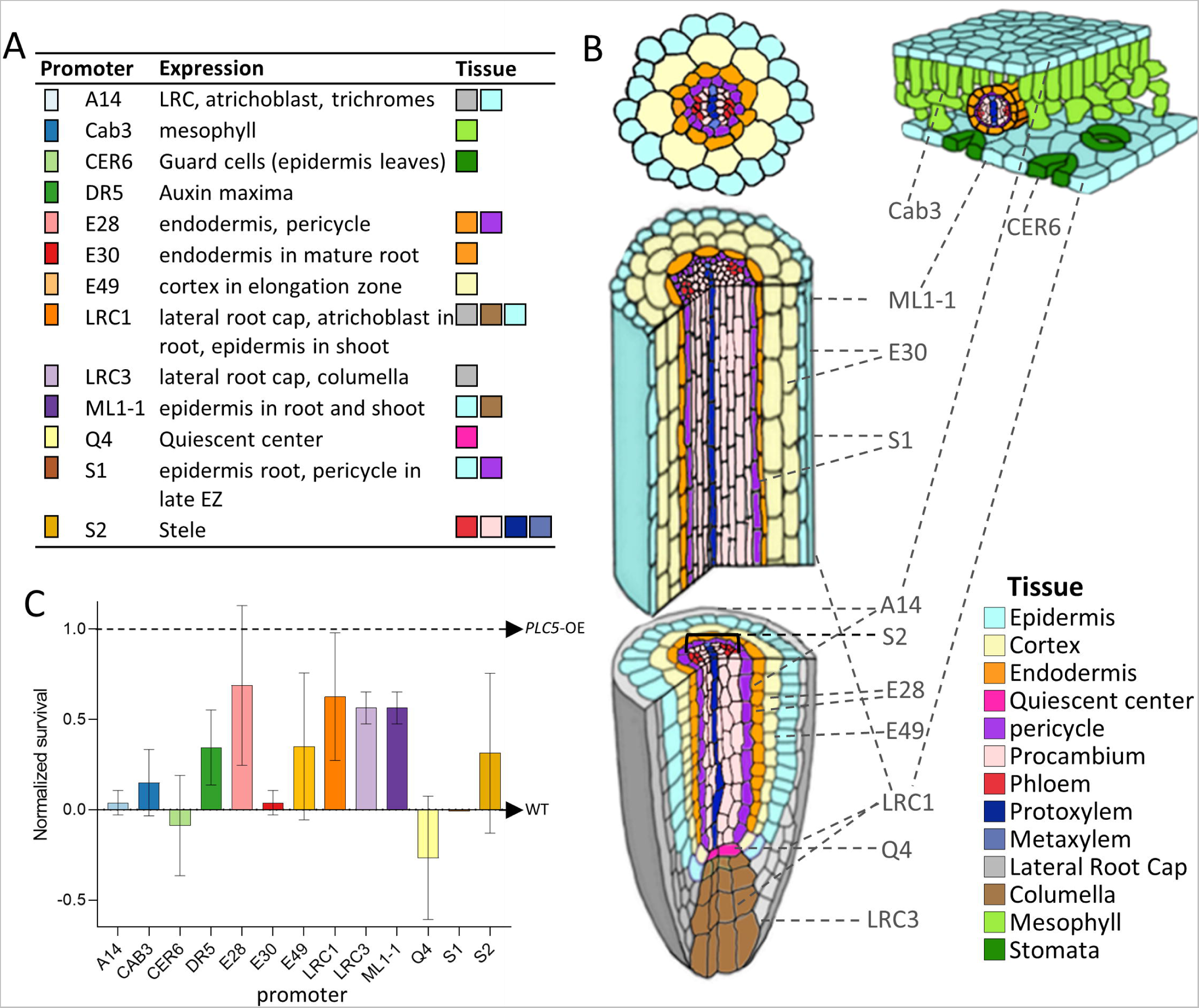
Survival under drought stress and localisation of *Tissue Specific Expression of PLC5*. A) Overview of promoter lines, and in which tissue they are mainly expressed. B) Schematic overview of in which tissue each line is expressed. Figure is adapted from (Kim *et al.,* 2017) and (Vaughan-Hirsch *et al.,* 2018) C) Plant survival of the *TSEP* lines after water stress. Three weeks-old plants were withheld from water for three weeks, after which they were watered again. Data was normalized using included WT as 0 and using the survival of the PLC5 OE line as 1. Each line was tested for 2 different inserts each tested 2 times over 3 different experiments.

Two independent transgenic lines for each construct were tested for their survival response to prolonged drought stress, with WT indicating the basal drought tolerance and UBQ10:*PLC5*-OE as a reference for the highest drought tolerance. As shown in Figure 8C, increased survival was obtained with six promoters, i.e. DR5, E28, E49, LRC1, LRC3 ML1-1. For the S2 promoter, only one insertion line showed the increased performance and was therefore excluded from subsequent analyses, as were *TSEP* lines that failed to show an increase in drought survival (i.e. A14, S1, CAB3 and CER6).

Interestingly, PCA plots of the six *TSEP* lines that showed enhanced drought survival were very similar to those of the *PLC-OE* lines (Fig. 9). At control conditions, no clear differences between WT and the *TSEP* lines were detected. However, *TSEP* lines did show distinct changes in response to water stress when compared to WT. The similarity in PCA plots between *PLC-OEs* and *TSEPs* indicated that similar phenotypes were occurring. Just like the *PLC-OE*s, *TSEP* lines revealed no significant difference in *Projected plant surface area* after normalization (Fig. 10A), indicating a similar response to water deprivation. At control conditions, however, these *TSEP* lines were significantly bigger than WT plants, with *pDR5:PLC5* plants showing the most prominent increase in size (Supplemental Fig. S20A), though this was lost again during water deprivation. *pLRC1:PLC5* plants were even significantly smaller in size at the final stage of water deprivation (Supplemental Fig. S20B).

**Figure 9.**
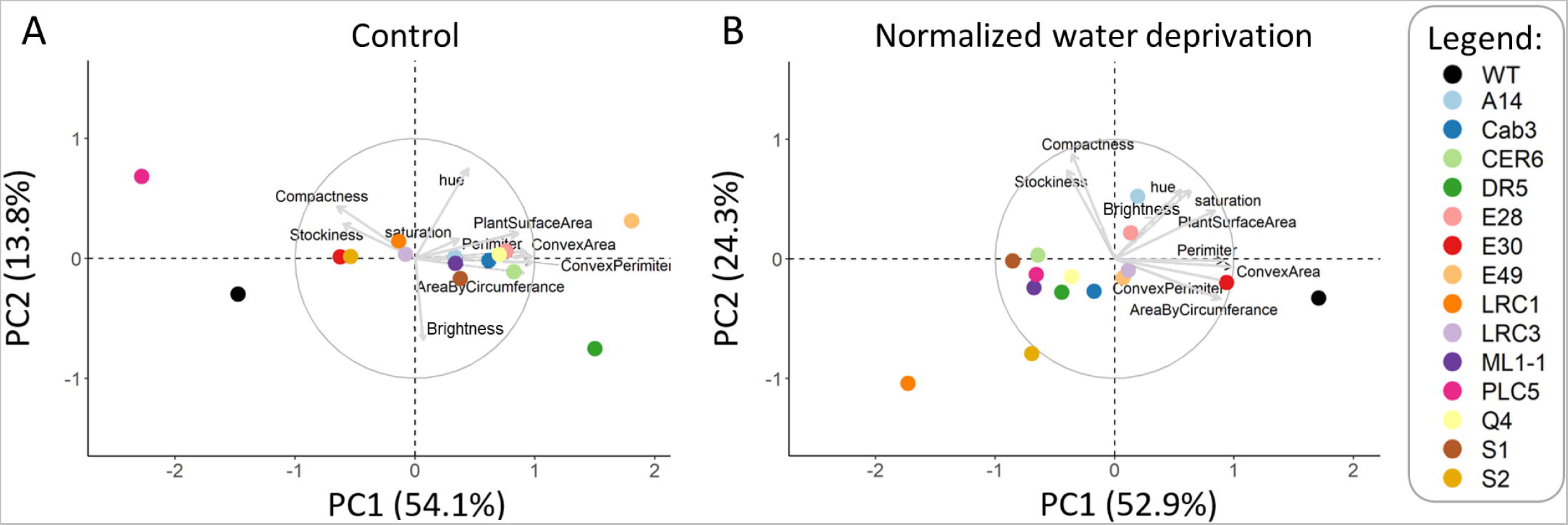
Altered plant morphology of *TSEP* lines compared to WT in reaction to water stress. PCA plots from control- (left panel, 21-45 DAS) and water-limiting conditions (right panel; 30-38 DAS (day 5-13 of stress). Data was normalized per day and per trait, to account for plants being larger at later days. Arrows indicate how much each trait contributes to each principal component (PC). Percentages of total variance represented by PC1 and PC2 are shown in parentheses. N = 40-60

**Figure 10.**
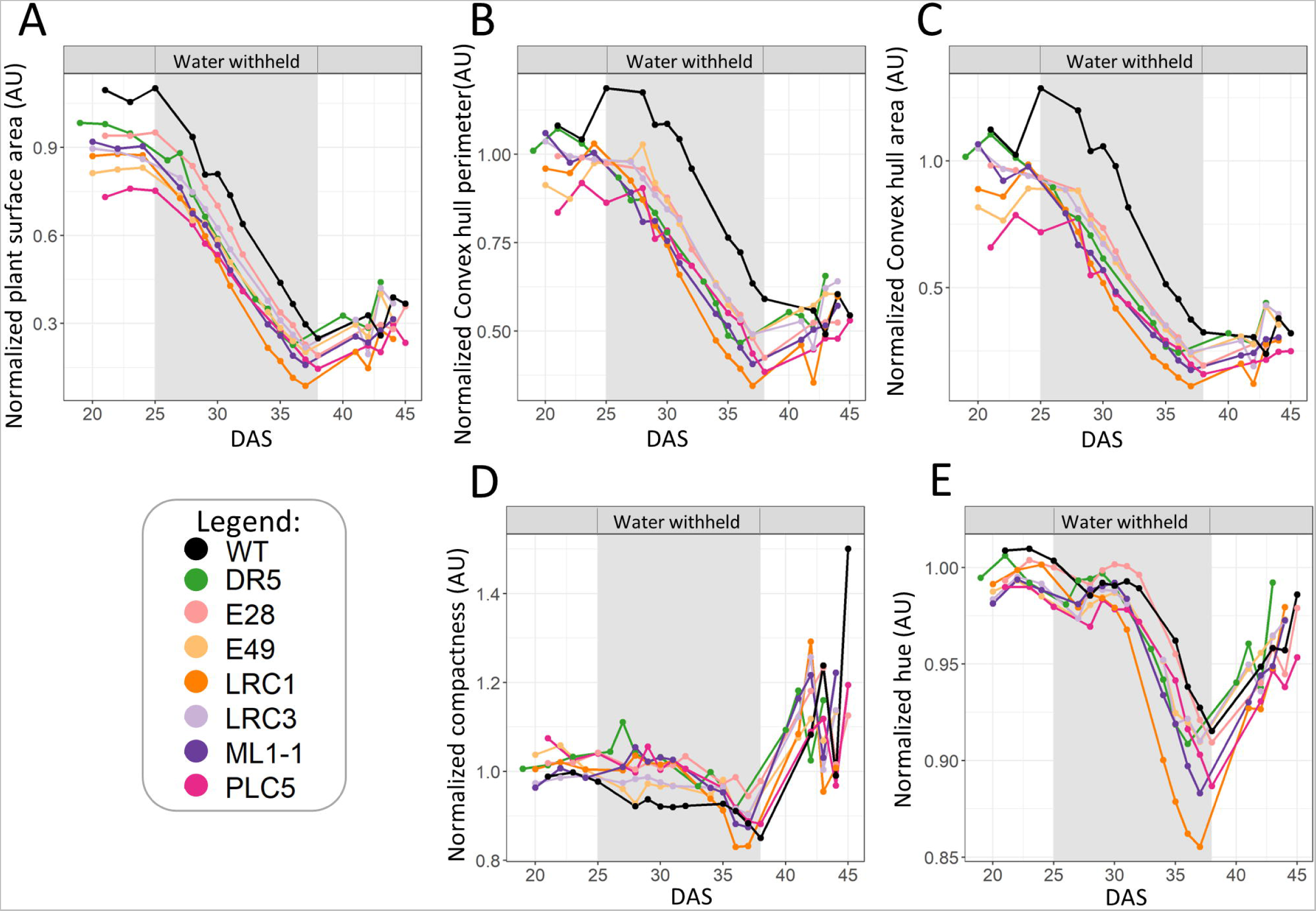
*TSEP* lines with increased drought survival respond similar to *PLC-OE* plants under water deprivation. All data is from water stressed plants normalized to control conditions. A) Projected plant surface area, B) Convex hull perimeter, C) Convex hull area, D) compactness and E) Hue.

The stronger decrease in *convex hull perimeter* and -*area* between WT and *PLC-OE* during water deprivation was again apparent for the *TSEP* lines (Figs. 10C & 10D), being statistically significant for the complete period of water-deprivation, for all lines. Similarly, the corresponding *compactness* was maintained during water stress for the *TSEPs*, only becoming less compact at the final days of water deprivation (Fig. 10D). *Hue* was not affected in most of the *PLC-OE*s under any of the experimental conditions, except for *PLC5*-OE, which showed higher green values. *TSEP* lines #E49 and #ML1-1 also revealed this increase in *Hue* at control conditions, as did #LRC1 and #LRC3 at later timepoints (Supplemental Fig. S16F). During water deprivation, the #LRC1 lines exhibited significantly lower *hue* (Supplemental Fig. S16G), and when the data were normalized, both #LRC1- and #ML1-1 lines responded significantly stronger than WT at later time points (Fig. 10E).

All six *TSEP* lines that exhibited increased drought survival (Fig. 8C) had increased biomass compared to WT under control conditions, in both FW and DW (Supplemental Figs. S21A, S121D). Under water deprivation, lines #DR5, #E28, #E49 and #ML1-1 were still significantly heavier than WT for both FW and DW. After normalization of the results of the water-deprived individuals to the ones of the controls, the differences in FW, DW, and RWC among the tested lines were no longer apparent. This indicates that all the lines had the same relative biomass penalty after two weeks of water deprivation followed by a week of rewatering.

## DISCUSSION

Earlier studies have shown that overexpression of *PLC* results in increased survival rates in response to water deprivation in various plant species (Wang *et al*., 2008; Georges *et al*., 2009; Tripathy *et al*., 2012), including Arabidopsis *PLC3, -4, -5*, and *7* (van Wijk *et al*., 2018; Zhang *et al*., 2018a; Zhang *et al*., 2018b; van Hooren *et al*., 2023). Here, we show that overexpression of *PLC2* and *PLC9* confirmed these results, suggesting that overexpression of <u>any</u> *PLC* can achieve this. While this is an interesting observation, it remains unclear whether plants also use endogenous PLC signalling as a protective reaction against water stress. A number of *PLC*s are indeed induced in response to drought, osmotic stress and or salinity (Hunt *et al.,* 2004; Tasma *et al.,* 2008; Li *et al.,* 2015) and PLC has also been linked to ABA, an important phytohormone in drought signalling (Hirayama *et al.,* 1995; Staxen *et al.,* 1999; Sanchez & Chua 2001; Hunt *et al.,* 2003; Mills *et al.,* 2004; Munnik 2014; Zhang *et al.,* 2018a).

To increase our understanding of how *PLC-OE* leads to increased drought tolerance, a large phenotyping experiment was initiated with two independent insertion lines of six distinct *PLC-OE*s, and 13 tissue-specific *PLC5*-expression lines. The PCA plot (Fig. 3 & Supplemental Fig. S1) clearly revealed that in response to water deprivation, *PLC-OE* lines grouped differently from WT. This was especially notable for *PLC9*-OE lines, as the PLC9 enzyme lacks two key histidines in the catalytic site (Ellis *et al.,* 1998). However, overexpression of *PLC9* clearly increased the survival rate under drought stress (Figs. 2B, 2C). Hence, we propose that either the change in the amino acids in the catalytic site does not interfere with the enzymatic function of PLC9, or that the increased survival found in *PLC9*-OEs is not directly linked to the activity of the enzyme. Other ways in which PLC9 could function is by forming protein-complexes through protein-protein interactions, or by preventing PPI signalling through shielding PPIs from binding other protein targets. Alternatively, if PLC enzymes would function as heterodimer with other PLCs, PLC9 could function as a silent partner.

That PLC9 has activity, directly or indirectly, is also indicated by earlier studies, showing that OE of *PLC9* increased heat stress tolerance, while *plc9* KO-mutants were more sensitive (Zheng *et al.,* 2012; Gao *et al.,* 2014; Ren *et al.,* 2017; Liu *et al.,* 2020). Another explanation for the observed mutations in the catalytic domain of PLC8 and PLC9, is that these members evolved an alternative catalytic domain. While two crucial histidines were lost, two new cationic amino acids were regained, i.e. E390K, and Y155R. Perhaps the later are used to accommodate the negative charge of the phosphorylated inositol headgroup. A possible advantage of the observed mutations could be that the enzymatic activity of PLC8- and PLC9 is less pH-sensitive, as Histidines lose their positive charge at pH>6, as opposed to Lysine and Arginine that maintain their charge at higher pH values. Heterologous PLC9 expression and *in vitro* enzyme activity assays will help investigating this further. Similarly, *PLC8-OE* lines could be generated to confirm the improved drought tolerance.

PCA plots revealed that *PLC5*-OE grouped differently from other *PLC-OE*s, which was particularly evident at control conditions. This is in line with earlier observations from Zhang *et al*. (2018b), where a ‘semi-dwarfed’ phenotype was documented. Dwarfism is often associated with drought tolerance, as small shoots transpire less water, and hence, more water remains available for the plant, making them appear as being more drought tolerant (Nishiyama *et al.,* 2011; Maggio *et al.,* 2018; Kudo *et al.,* 2019; Bhatnagar *et al.,* 2020; Yang *et al.,* 2020). However, none of the other *PLC-OE* lines exhibited this ‘semi-dwarfed’ phenotype, so the reason for the increased drought tolerance cannot be simply due to dwarfism. Moreover, at control conditions, the rest of the *PLC-OE* lines tend to grown bigger (Fig. 4A) and produce more biomass (Supplemental Figs. S19A &S19D). Under water deprivation, this difference became less significant, though it was still present in some of the tested lines (Fig. 4B; Supplemental Fig. S19B, S19E). When specifically looking at the rate of growth decline, also no change was observed with the exception of *PLC9*-OEs, which declined stronger (Fig. 4E). Normalization of the water-deprived plants showed that drought stress leads to a non-significant decrease in size (Fig. 4C), which was compensated after recovery (Supplemental Figs. S19C & S19F). Hence, we conclude that the shared increase in drought survival of the *PLC-OE*s is not due to a growth penalty.

Most stresses, including water stress, are associated with a decrease in photosynthetic activity, and eventual decline in the amount of chlorophyll (Chaves *et al.,* 2009). In this experiment, we saw the same thing, but found no difference between WT and OE lines in *F_v_/F_m_* ratios (Supplemental Fig. S6), nor for *Hue*, a readout for chlorophyll content (Supplemental Fig. S16). At control conditions, *PLC5*-OE lines were significantly greener than WT, though increased chlorophyll levels may be explained by the reduced cell size, which could also be the reason for the smaller shoot phenotype. This indicates that the increased survival in the OE lines was not caused by changes in photosynthetic capacity, though PLC interference leading to increased ROS turnover and an enhanced stress response, could still be a possibility. It should be noted that *F_m_* level, which indicates the amount of active reaction centres available for photosynthesis, was higher in all the *PLC-OE* lines at control conditions, though this was not statistically significant (Supplemental Fig. S5A).

The PCA plots did reveal changes correlating to the shoot architecture. In response to water deprivation, both C*onvex hull -perimete*r & -*area* decreased; an effect that was significantly greater in all *PLC- OE*s. Since *PLC-OE* lines tend to be bigger than WT, and hence are expected to have larger perimeters, the two factors appear to be correlated. However, when plotting the two parameters against each other, a smaller perimeter and area for the *PLC-OE*s were recorded when plants of the same size were compared, indicating that a change in the architecture has occurred (Figs. 5C, 5E, 6C and 6E). The ratio of the *Projected plant surface area* and *Convex hull area* can be given as a *Compactness* value, and under water deprived conditions, the *Compactness* of the *PLC-OE*s as a group, was significantly more affected than for WT (Fig. 7D).

No effect in *Relative water content* at any condition for any of the tested lines, as compared to WT was found (Supplemental figs. S19G, S19H, S21G, S21H), so it is unlikely that *PLC-OE* has a beneficial effect for the plants giving them the advantage to deal better with low water potential (which is linked to the drought tolerance strategy). We do find that *PLC-OE* results in a stronger response related to shoot size and architecture, traits that are more associated with an increased drought-avoidance strategy.

The increased survival under water deprivation is a common characteristic feature of all tested, independent *PLC-OE* lines. In this study, an architectural change in response to water stress was found. This modification was not limited to the *PLC-OE* lines but was also found in the six *TSEP* lines that exhibited increased survival under water deprivation (Figs. 8 & 10). The fact that the architectural changes and increased-survival phenotypes can be reproduced by expressing *PLC5* in various tissues, indicates that the phenotypes from *PLC- OE*s are probably caused by ectopic expression and not from a higher expression in tissues where *PLC* is already expressed.

Of the 13 *TSEP* lines constructed, six showed an increased survival phenotype in response to water deprivation in two independent insertion lines. The expression of these promoters is mainly restricted to the developing parts of the root. One of them, #E28 targets the expression of the transgene in the root pericycle and endodermis, while #E49 is specifically expressed in the cortex of the elongation zone. In contrast, #E30 line which did not show increased survival to water deprivation, targeted the expression of *PLC5* in the endodermis of the root maturation zone. #S1, similarly, gives expression in the root epidermis and pericycle, but only in the late elongation zone. The different tissues in the mature root have important physiological role, but it appears that the targeted *PLC* expression therein does not affect the survival rate under drought stress. Both promoters #LRC3 and #LRC1 are mostly active in the Lateral Root Cap (LRC), but #LRC1 also triggers the expression in the root and shoot epidermis, similarly to #ML1-1. The LRC plays an important role in the early development of the root (Di Mambro *et al.,* 2019). It is the primary anatomical element to get in contact with the surrounding environment, thus it is the first to sense, and probably prone to react to, the changes in parameters like temperature, water and nutrients availability or mechanical barriers.

#DR5 expression coincides with auxin maxima (Ulmasov *et al*., 1997), and thus, in the case of roots, with the zones of intensive growth. Consequently, it has a high overlap with the sites of expression of all the other constructs that target *PLC5* expression in developing parts of the root. The absence of any effects on the survival rates of the #CAB3, #CER6 and #Q4 transgenic lines indicates that the ectopic expression of *PLC5* in either mesophyll cells, guard cells or the quiescent centre alone does not provide physiological advantages for better survival under water deprivation. Most of the parameters evaluated in the present study are associated with the above ground parts of Arabidopsis, however, the observed pattern of increased survival of lines with targeted *PLC5* expression in the most dynamic parts of the root (cell division and elongation zone), gives a new organ-specific direction for further research on the role of PLCs in drought perception and/or its response.

Even though there were rosette size differences in the six *TSEP* lines that exhibited increased survival, with #DR5 being by far the largest and #LRC1 plants having the smallest rosettes among all water-deprived individuals, they all appeared more compact under water stress. While this parameter can easily be measured, in big phenotyping experiments, it is often ignored, with a few exceptions (Skirycz *et al.,* 2011; Prerostova *et al.,* 2018; Marin-de la Rosa *et al.,* 2019). Specific studies on compactness in Arabidopsis revealed that it generally decreases as plants grow older (Marin-de la Rosa *et al.,* 2019). A circadian rhythm on compactness has also been reported, which most likely reflects the change in leaf angle during day and night, being almost horizontal, and therefore the least compact during midday, and becoming more compact at night (Dhondt *et al.,* 2014). This is unlikely to ultimately affect the data presented in this study, as plants were randomly measured.

In response to salt stress or H_2_O_2_, though not in response to mannitol, plants tend to become more compact (Jansen *et al.,* 2009; Dhondt *et al.,* 2014). A slight compactness decrease has been found in response to drought Marin-de la Rosa *et al.,* 2019). Our data confirmed that the compactness of plants decreased while growing older, and that water-limiting conditions induced an even stronger decrease. Since compactness is easily measured, and in automated phenotyping experiments often is, and is affected by (abiotic) stress, a greater focus on this trait could help future research to characterise and identify stress responses even better.

An increase in *compactness* could result from different variables, including leaf shape, leaf angle, and shorter petiole length. However, no significant difference in *leaf diameter* measurements was found among all tested lines (Supplemental Fig. S28), and *PLC-OE*s plants even trend to exhibit narrower leaves, arguing against the idea that leaf shape is the main cause for the observed differences in compactness between the WT and the *PLC-OE*s under water deprivation.

Low turgor pressure may cause leaves to wilt, thus decreasing their standing angle and subsequently the *Compactness* of the monitored individual. However, if this would have been the sole explanation of the detected difference in *Compactness,* then we would expect the alteration to occur when plants stop growing. However, the difference was already observed within the first few days of water deprivation.

Changes in petiole elongation have not been well studied in the context of drought stress. Most published data is related to its role in thermomorphogenesis, where plants grow longer petioles and increase their hyponasty when grown at temperatures above 26-28 °C, which decreases their compactness substantially (Ibanez *et al.,* 2017). One of the main mechanisms by which plants are able to cool down under heat stress is to evaporate water (transpiration). This process is limited by the water supply and air humidity. Modelling of the air flow around heat-stressed leaves predicts that a decrease in compactness would increase the air flow around the leaves, improving the removal of water-saturated air around the leaves, and enabling a steady supply of air to evaporate water into (Bridge *et al.,* 2013). This finding is not surprising, as transpiration is the exact same mechanism in which many mammals cool down. If increasing the petiole length can facilitate water evaporation, then the opposite, an increase in compactness, might help in conserving water. A compact canopy would reduce air flow underneath the leaves, making the air therein increasingly humid. Water evaporates badly in humid air, therefore the stomata that remain open at the same level would lose less water (Gonzalez *et al.,* 2012).

So far, research on *Compactness* has mainly centred on grasses (grains and rice), in order to grow more plants in the same space. Interestingly, also in grasses an increase in drought tolerance has been correlated with *Compactness* (Honsdorf *et al.,* 2014; Kim *et al.,* 2020; Gano *et al.,* 2021). Hence, this correlation might be interesting for breeding crops as well, as it simultaneously invests into drought tolerance based on drought avoidance rather than drought-escape. Further research into this correlation may further substantiate the ideas brought forward here. The next challenge will be to unravel the molecular mechanism behind this phenomenon.

## MATERIAL & METHODS

### Plant material

*Arabidopsis thaliana* ecotype Col-0 was used as genetic background for all the lines used in this research. Plants were grown for at least one generation together and synchronised seed batches were used. In the GROWSCREEN-FLUORO setup, the means of two Col-0 wild types (WTs) were used: one being the line that was grown together with the *PLC-OE* lines for several generations, and the other was freshly ordered from NASC (Scholl *et al.,* 2000). UBQ10:*PLC3* (AT4G38530) #9 & #16, UBQ10:*PLC5* (AT5G58690) #2 & #3, and UBQ10:*PLC7* (AT3G55940) #9 & #17were described earlier (van Wijk *et al.,* 2018; Zhang *et al.,* 2018a,b). The 35S:*PLC4* (AT5G58700) #2 & #4 line was kindly provided by Dr. Rodrigo Gutiérrez (Universidad Catolica de Chile). *PLC2-*OE (AT3G08510) *PLC9*-OE (AT3G47220) and 13 *TSEP* lines were generated in this study and will be described below. KO *plc9-1* (SALK_025949) and *plc9-2* (SALK_021982) lines were obtained from SALK (signal.salk.edu) and a *plc2- 1* mutant in Ws backgound was kindly provided by Dr. Ana Laxalt (University of Mar del Plata, Aregentina; Di Fino *et al.,* 2017).

### Plant growth conditions

For genotyping, qPCR and western-blot analysis, plant material was grown on agar plates. Seeds were first surface sterilized in a 3.5 L desiccator with 20 ml of thin bleach and 600 µl of 37% w/w HCl for 3-4h, and then stratified in 0.1% (w/v) daishin agar at 4°C for 2-4 days in the dark. Stratified seeds were sown on square Petri dishes (Greiner) containing 40 ml of ½ MS medium, supplemented with 1% (w/v) sucrose, 0.1% (w/v) MES (pH 5.8) and 1% (w/v) daishin agar (Duchefa Biochemie). Plates were placed vertically at an angle of 70° at long-day conditions (22°C, RH 70%, 16h of light, 8h of dark), and scanned at different time points (Epson Perfection V700).

### Construction of PLC2-OE, PLC9-OE and TSEP lines

*pUBQ10:PLC2, p2x35SP:PLC2, pUBQ10*:*PLC9* and 13 ’*Tissue-Specific Expression of PLC5’* (*TSEP*) lines were constructed using gateway cloning. Oligonucleotide primers including Atb1 and AtB2 sites were used to PCR amplify the CDS of *PLC2* and *PLC9* from cDNA (Supplemental Table S1). The PCR fragment was inserted into the pDONR207 vector with BPII clonase enzyme mix to generate a BOX2-vector entry clone and entry clones for pGEM-UBQ10, pGEM-2x35S and pGEM-StopTnos vectors were kindly provided by Dr. Ben Scheres (Utrecht University, The Netherlands). The three entry clones were recombined in a pGreenII-125 expression vector using the Multisite Gateway Three-Fragment Construction Kit (www.lifetechnologies.com) and the supplier’s protocol, and checked by PCR and subsequent sequencing. (https://www.thermofisher.com/nl/en/home/life-science/cloning/gateway-cloning/gateway-technology.html).

TSEP lines were generated by recombining the tissue-specific promoters (Supplemental Table 2; (Vaseva *et al.,* 2018) in pENTR5 vectors, the coding sequence of *PLC5* in pGreenII029JV (Zhang *et al.,* 2018b), and pGEM-StopTnos in a pGreen0125-expression vector using Multisite Gateway Cloning. Constructs were validated for *PLC5* mutations by PCR and subsequent sequencing. *PLC5* was chosen since pilot studies indicated *PLC5*-OE had the strongest increase in drought survival.

All constructs were co-transformed with vir-helper vector, pSOUP into *Agrobacterium tumefaciens* strain GV3101, which was then used to transform Arabidopsis by floral dipping (Clough & Bent 1998). At least two homozygous lines were selected by norflurazon selection in T3 generation, and used in the next experiments. The norflurazon selection was used as a proxy to determine successful integration and expression of the *TSEP* constructs.

### RNA extraction and qPCR

*PLC2-* and *PLC9* expression in KO- and OE lines was analysed using qPCR. Total RNA was extracted from 7-day old seedlings that were homogenized in liquid nitrogen with mortar and pestle using a homemade Trizol reagent. RNA (0.5 μg) was converted into cDNA using oligo-dT18 primers, dNTPs, and SuperScript III Reverse Transcriptase (Invitrogen), according to the manufacturer’s instructions. Q-PCR was performed using HOT FIREPol EvaGreen qPCR mix Plus (ROX) (Solis Biodyne) with an ABI 7500 Real-Time PCR System (Applied Biosystems). Relative expression levels were determined by comparing the threshold cycle values for *PLC2* (AT3G08510) or *PLC9* (AT3G47220) to the reference gene, *SAND* (AT2G28390) (Czechowski *et al.,* 2005). Primers for *SAND, PLC2* and *PLC9* (being negative for *PLC8*) are listed in Supplemental Table S1. Three technical replicates and three biological replicates were used, with each biological replicate consisting of 12-15 seedlings. Changes were tested for significance with a Student T-test.

### Protein isolation and Western-blotting

For protein extraction, 50 µl of ice-cold protein extraction buffer (0.5% (v/v) NP-40, 75 mM NaCl, 100 mM Tris-HCl (pH 8.0), 1% (w/v) PVPP, 100 mM DTT, 1 mM PMSF, 10 mM EDTA (pH 8.0), supplemented with protease inhibitor EDTA free (Roche) just before use) was added to 10 mg root material of 5 DAG seedlings grown on plates that was grinded in liquid nitrogen. During the whole procedure, material was mixed well and kept cold. Samples were centrifuged 2x 5 min at 2000g followed by 2x 5 min at 5000g; each time transferring the supernatant to a new tube. Protein concentrations were estimated using Nanodrop (ThermoFisher).

SDS-PAGE gels (10%) were loaded with 4-5 µg of boiled and centrifuged protein sample (10 min 95°C; 2 min 16.000g) in sample buffer (containing 2% SDS (w/v), 10% glycerol (v/v), 5% ß-mercaptoethanol (v/v), 60 mM Tris-HCl pH 6.8, 0.02% (w/v) bromophenol blue), and ran for ∼1h at 30 mA/gel. Gels were blotted to PVDF membrane (Immobilon-P, Merck) that were then incubated for 1 hr with a PLC2-antibody (1:2000) provided by Dr. Ana Laxalt (D’Ambrosio *et al.,* 2017). Filters were washed 4 times for 5 min with a phosphate-buffered saline (PBS) buffer containing 0.05% Tween-20 (v/v), incubated with a second antibody, polyclonal Goat anti-rabbit IgG (H+L)/HRP (1:5000; ThermoFisher) for 1 hr, washed 6x 10 min with PBS buffer containing 0.05% Tween-20 (v/v), and imaged using ECL.

### Drought tolerance assay

Seeds were stratified in liquid for two nights in the dark at 4 °C, and thereafter directly sown in soil (pots with diameter from 3.8 to 5 cm and height of 5 cm for *PLC9-OE*, and pots of 4.5 x 4.5 cm and height of 7.5 cm for *PLC2-OE*), containing 30 g or 80 g of soil, respectively (Zaaigrond nr. 1, SIR 27010-15, JongKind BV, The Netherlands). Plants were grown at short-day conditions (22 °C with 13 h light/11 h darkness).

In the drought experiment with *PLC9-OE* and *TSEP* lines, 15 plants per line were grown for 3 weeks and watered every other day. After these 3 weeks, the excess of water was removed from the trays, and the plants were withheld from water for 3 weeks. At the end of the experiment, plants were rewatered again to allow the survived individuals to recover and turn green. During the experiment, pots were randomized twice per week, using a shuffling schedule that would distribute the plants equally on random positions at the in- and outside of a tray and in the corners. The experiment was performed 3 times, with each line included at least 2 times. In the drought experiment with *PLC2-OEs*, nine plants per pot were grown for 37 days in well-watered conditions, after which plants were left to dry for an additional 18 days, with pots being randomized twice a week. In both experiments, plants were re-watered again after the drought period and left to recover for nine days before pictures were taken. The experiment was performed twice, giving similar results.

### PLC9 protein structure modelling

The crystal structure *PLC-δ1 from* of *Rattus norvegicus (1DJX)* (Essen et al., 1997) was used as a template to build models of the Arabidopsis PLC5- and PLC9 proteins using https://evcouplings.org/, which were then further visualized using Novavold® (Version 17.2; DNASTAR, Madison, WI).

### Phenotyping platform and drought stress setup

For the phenotyping, the GROWSCREEN-FLUORO setups in the GROWSCREEN chamber facilities of IBG-2 at Plant Sciences of the Forschungszentrum Jülich (Jülich, Germany) were used, as has been described earlier (Jansen *et al.,* 2009; Barboza-Barquero *et al.,* 2015; `Prerostova *et al.,* 2018). Single seeds were sown in multi-well trays filled with a clay-peat substrate (Einheitserde Typ Mini-Tray, Einheitserdewerke Werkverband e.V., Sinntal- Altengronau, Germany) using a *pheno*Seeder robot (Jahnke *et al.,* 2016). After stratification for four days at 4 °C, trays were transferred to a climate-controlled GROWSCREEN chamber at 22 °C/18 °C (day/night), 50% relative air humidity and an 8/16 h light/dark period. Seed germination and seedling growth was monitored daily until transplantation using an automated imaging and image processing system. Fourteen days after sowing, seedlings with the largest leaf surface area per line, determined by an automated system, were transplanted to pots (7 × 7 × 8 cm) filled with peat-sand-pumice substrate (SoMi 513 Dachstauden; Hawita, Vechta, Germany), and watered to field capacity. Pots were arranged in trays that were positioned in two independent GROWSCREEN chambers, set at the same environmental conditions as for germination and seedling growth. Pots were first dried to 60% to 40% of field capacity and kept at that moisture level until the start drought treatment (withholding water) at 25 days after sowing (DAS). Pots of the control treatment were maintained between 60 and 40% of field capacity until harvest at 45 DAS, whereas pots of the drought treatment were dried to between 8% and 5% of field capacity over a period of 13 days. Pots were then re-watered to 60% of field capacity and left to recover for seven days until harvest at 45 DAS. Shoot biomass was measured at harvest as fresh weight, followed by dry weight, after drying rosettes for seven days at 65 °C.

Per construct, two independent insertion lines were used, and for each insertion line (and WT), 30 plants were used per treatment, making 2460 plants in total. All plants were distributed according to a complete randomized block design across a total of 66 trays with 40 pots each. Trays for both treatments were placed in the GROWSCREEN chambers where a robot daily delivered the trays to a weighing station and randomized them by placing them back in another position. Trays were watered twice per week. The robot also delivered the trays to the GROWSCREEN-FLUORO imaging station. Imaging started at 21 DAS and was repeated every 2-3 days until harvest for red/green/blue values (RGB) imaging. From 28 to 38 DAS, during the water-withholding period, RGB imaging was performed daily. Chlorophyll fluorescence of dark-adapted plants was imaged twice per week. Image analysis was performed by in-house developed software and included the RGB images, a first rough segmentation of shoots from the substrate background using thresholds on *Hue*, *Saturation* and *Brightness* values. This was followed by a precise segmentation based on a trained support-vector machine algorithm. The obtained phenotypic traits included *projected shoot surface area*, *shoot perimeter*, *convex hull area*, *convex hull perimeter*, *compactness*, *surface coverage*, *stockiness*, *area by circumference*, and the average *red*, *green* and *blue* values, and *hue*, *saturation* and *brightness* values of all shoot pixels. Chlorophyll fluorescence data were processed as described in Jansen et al. (2009), resulting in the traits *F_0_*, *F_m_* and *F_v_/F_m_*.

### Data processing

Data analysis was performed in R, using a script shared at https://github.com/MaxvHooren/GROWSCREEN-analysis. Because the plants from one of the climate chambers were slightly bigger, a correction was applied, which was determined by calculating the difference between the chambers per day and per treatment. These differences were averaged and then applied as a correction factor. This process was repeated for every trait measured. Principal component analysis (PCA) was performed with the FactoMineR and FactoExtra packages. The data was normalized per day before doing PCA analysis, as older/bigger plants would have had a bigger impact on the estimated differences.

As not all seeds were sown on the same day, measurements and DAS were not aligned. Therefore, it was difficult to establish the relative growth rates per day and to compare the changes in this parameter between different groups, as different numbers of plants were measured per line on each day. In order to overcome this, growth curves were fitted to ‘rosette projected-surface area’ over time for each individual plant to enable the calculation of growth rate. For data obtained during control conditions, an exponential growth curve (y=a*e^Bx^) was fitted, whereas a quadratic function (y=ax^2^+bx+c) was used for data obtained during water withholding.

The statistical significance among tested lines for every day was determined by ANOVA test, followed by a TUKEY HSD test, comparing the values to the combined WT values (Supplemental Table S4). Blank timepoints correspond to times when either the line in question or WT was not measured.

After the analysis in R, bar graphs were built using GraphPad Prism version 8.4.2 for Windows (GraphPad Software, San Diego, CA, USA; www.graphpad.com).

## DATA AVAILABILITY STATEMENT

The raw data used in this paper can be found at the GrowSreenChamber institution on the “Phenomis” database at IBG-2. The script for analysis can be found at https://github.com/MaxvHooren/GROWSCREEN-analysis. Analysed data is available at https://zenodo.org/record/7715023. All other data supporting the findings of this study are available from the corresponding author (TM)

## ACKNOWLEDGEMENTS

We thank Rodrigo Gutiérrez (Universidad Catolica de Chile) for the *PLC4-OE* lines. We thank Fabio Fiorani, Silvia Braun and Nathalie Wuyts from IBG-2 at Plant Sciences of the Forschungszentrum Jülich (Jülich, Germany) for help with the experimental design, conducting the GROWSCREEN experiment, and reviewing the manuscript.

## AUTHOR CONTRIBUTIONS

MvH and TM designed the experiments. RvW created the *PLC2-OE* and *PLC9-OE* lines. MvH created the tissue specific *PLC5-OE* (*TSEP*) lines using the promoter vectors created by IV and DvdS. RvW did qPCR, western blot analysis and drought experiments on *PLC2-OE*. MvH did qPCR and drought experiments on *PLC9-OE* and *TSEP* lines. MvH helped with sowing, transfer, and harvesting of the GROWSCREEN experiment, which itself was conducted by IBG-2 at Plant Sciences of the Forschungszentrum Jülich. MvH performed the data analysis and MvH and TM constructed the figures and wrote the article. RvW, IV, DvdS and MH critically read the ms and made valuable corrections.

## SUPPLEMENTARY INFORMATION

Supplemental Table S1. Primers for the construction of PLC9 OE and tor the PLC9 qPCR

Supplemental Table S2. Promoters used for Tissue Specific Expression of PLC5

Supplemental Table S3. Key amino acids of the catalytic site of rat PLCδ1 and their counterparts in Arabidopsis PLCs

Supplemental table 4 - Significance of traits in phenomics experiment

Supplemental Figure 1. PCA plot of Chlorophyll fluorescence camera traits.

Supplemental Figure 2. Darkyield of plants at control- and water limiting conditions.

Supplemental Figure 3. Eccentricity of plants at control- and water limiting conditions.

Supplemental Figure 4. F0 of plants at control- and water limiting conditions.

Supplemental Figure 5. Fm of plants at control- and water limiting conditions

Supplemental Figure 6. Fv /Fm ratios of plants at control- and water limiting conditions.

Supplemental Figure 7. Insidediameter of plants at control- and water limiting

Supplemental Figure 8. Midyieldfluoresence of plants at control- and water limiting conditions.

Supplemental Figure 9. Minimal leaf diameter of plants at control- and water limiting conditions.

Supplemental Figure 10. Outside leaf diameter of plants at control- and water limiting conditions.

Supplemental Figure 11. Average leaf diameter of plants at control- and water limiting conditions.

Supplemental Figure 12. Number of leaves of plants at control- and water limiting conditions.

Supplemental figure 13. Plant perimeter at control- and water limiting conditions.

Supplemental figure 14. Area by circumference at control- and water limiting conditions

Supplemental figure 15. Stockiness at control- and water limiting conditions.

Supplemental figure 16. Hue at control- and water limiting conditions.

Supplemental figure 17. Brightness at control- and water limiting conditions.

Supplemental figure 18. Saturation at control- and water limiting conditions

Supplemental figure 19. Fresh- and dry weight and relative water content in WT and PLC-OE lines.

Supplemental figure 20. Projected plant surface area of TSEP lines under control and water limiting conditions.

Supplemental figure 21. Fresh- and dry weight and relative water content in WT and TSEP lines.

Supplemental figure 22. Convex hull permiter at control- and water limiting conditions

Supplemental figure 23. Convex hull area at control- and water limiting conditions.

Supplemental figure 24. Compactness at control- and water limiting conditions.

Supplemental figure 25. Surface coverage at control- and water limiting conditions.

Supplemental figure 26. RGB Blue values at control- and water limiting conditions

Supplemental figure 27. RGB Green values at control- and water limiting conditions

Supplemental figure 28. RGB Red values at control- and water limiting conditions

