## Supplemental table 1 -3 and figures 1- 28 for "Ectopic Expression of Distinct *PLC* Genes Identifies ‘Compactness’ as Novel Architectural Shoot Strategy to Cope with Drought Stress"

**Supplemental Table S1**. Primers for the construction of PLC9 OE and tor the *PLC9* qPCR

| **Primers** | **Sequence** |
|---|---|
| PLC9_fwd | ggggacaagtttgtacaaaaaagcaggctATGGTGAATTTAAGAAAGAAGTTTG |
| PLC9_rev | ggggaccactttgtacaagaaagctgggtTCTAAGACCACTTAAAACGTGTGAGC |
| attB1-PLC2fw | ggggacaagtttgtacaaaaaagcaggctATGTCGAAGCAAACGTAC |
| attB2-PLC2rv | ggggaccactttgtacaagaaagctgggtCACAAACTCCACCTTCACGA |
| LB1 | GCCTTTTCAGAAATGGATAAATAGCCTTGCTTCC |
| PLC9_qPCR_fwd | ATCGTGACATGAATGCTCCA |
| PLC9_qPCR_rev | ACGCATATGCCGTCTTTACC |
| Q_PLC2_set1_fw | TTTGGAGGATTTGTATGTGTGAGAG |
| Q_PLC2_set1_rv | CCACAAATCATCATACGATTCAAGTG |
| SAND_qPCR_fwd | CAGACAAGGCGATGGCGATA |
| SAND_qPCR_rev | GCTTTCTCTCAAGGGTTTCTGGGT |

**Supplemental Table S2**. Promoters used for Tissue Specific Expression of *PLC5*

| Promotor | Expression | Corresponding gene | |
|---|---|---|---|
| A14 | LRC, atrichoblast, trichromes | AT5G43040 | KCS6 |
| Cab3 | Mesophyll | AT1G29910 | LHCB1.2 |
| CER6 | Guard cells (epidermis leaves) | AT1G68530 | Cysteine/Histidine-rich C1 domain family protein |
| E28 | Endodermis, pericycle | AT4G00940 | DOF4.1 |
| E30 | Endodermis in mature root | AT4G21340 | B70 |
| E49 | Cortex in elongation zone | AT3G05150 | Major facilitator superfamily protein |
| LRC1 | Lateral root cap, atrichoblast in root, epidermis in shoot | AT1G08930 | ERD6 |
| LRC3 | Lateral root cap, columella | AT1G30650 | WRKY14 |
| ML1-1 | epidermis in root and shoot | AT4G21750 | ATML1 |
| Q4 | Quiescent centre | AT1G69490 | ANAC029 |
| S1 | Epidermis root, pericycle in late EZ | AT4G27410 | RD26 |
| S2 | Stele | AT3G61850 | DAG1 |
| DR5 | Auxin maxima | | |

**Supplemental Table S3.** Key amino acids of the catalytic site of rat PLCδ1 and their counterparts in Arabidopsis PLCs. Differences are given in red. Based on Ellis et al. (1998)

| | Catalytic site | | | | | | | | |
|---|---|---|---|---|---|---|---|---|---|
| RnPLCδ1 | H:311 | H:356 | Y:155 | E:341 | D:343 | E:390 | K:440 | S:522 | R:549 |
| AtPLC1 | H:120 | H:166 | F:352 | E:150 | D:152 | E:200 | K:249 | S | R |
| AtPLC2 | H121 | H:164 | Y:375 | E:148 | D:150 | E:198 | K:248 | S | R |
| AtPLC3 | H:118 | H:167 | Y:354 | E:151 | D:153 | E:201 | K:250 | S | R |
| AtPLC4 | H:129 | H:174 | Y:391 | E:159 | D:161 | E:208 | K:257 | S | R |
| AtPLC5 | H:127 | H:173 | Y:372 | E:157 | D:159 | E:207 | K:256 | S | R |
| AtPLC6 | H:152 | H:198 | Y:407 | E:182 | D:184 | E:232 | K:281 | S | R |
| AtPLC7 | H:118 | H:164 | Y:381 | E:148 | D:150 | E:198 | K:248 | S | R |
| AtPLC8 | L:120 | P:164 | R:327 | E:149 | D:151 | K:198 | R:247 | - | R |
| AtPLC9 | L:122 | P:168 | R:326 | E:153 | D:155 | K:203 | R:253 | - | R |

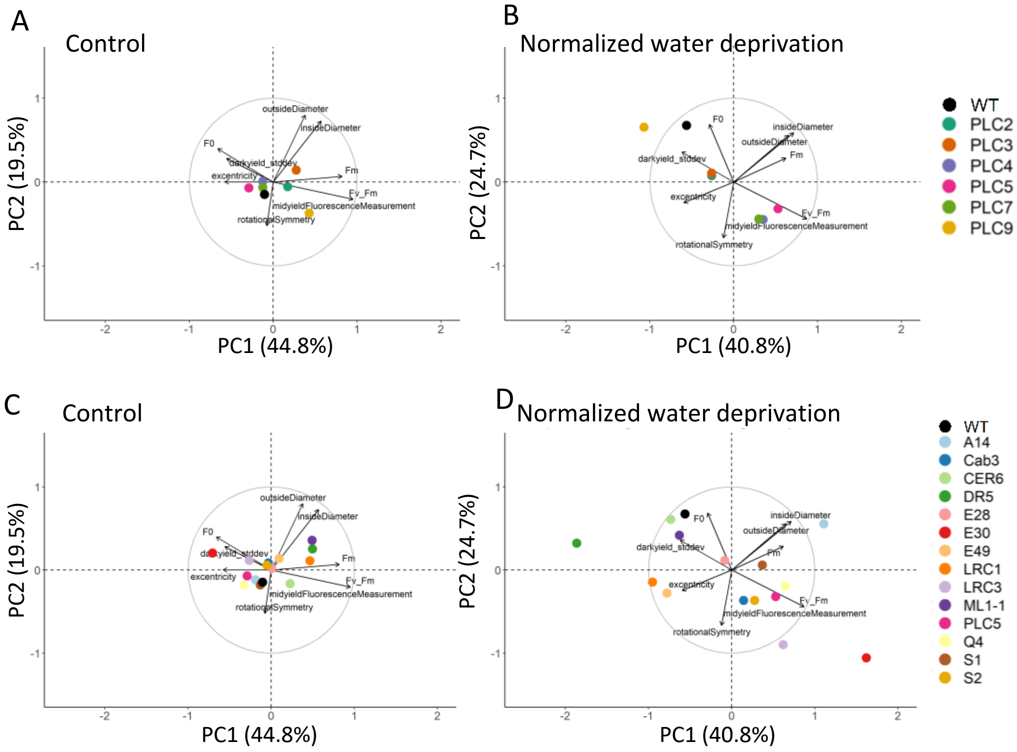


**Supplemental Figure 1**. PCA plot of Chlorophyl fluorescence camera traits.
PCA plots from control (panel A and C; 21-45 DAS) and water-limiting conditions normalized to control (panel B and D; 30-38 DAS (day 5-13 of stress). Data was normalized per day and per trait, to account for plants being larger at later days. Arrows size and direction indicate how much each trait contributes to each principal component (PC). Percentages of total variance represented by PC1 and PC2 are shown in parentheses. N = 40-60

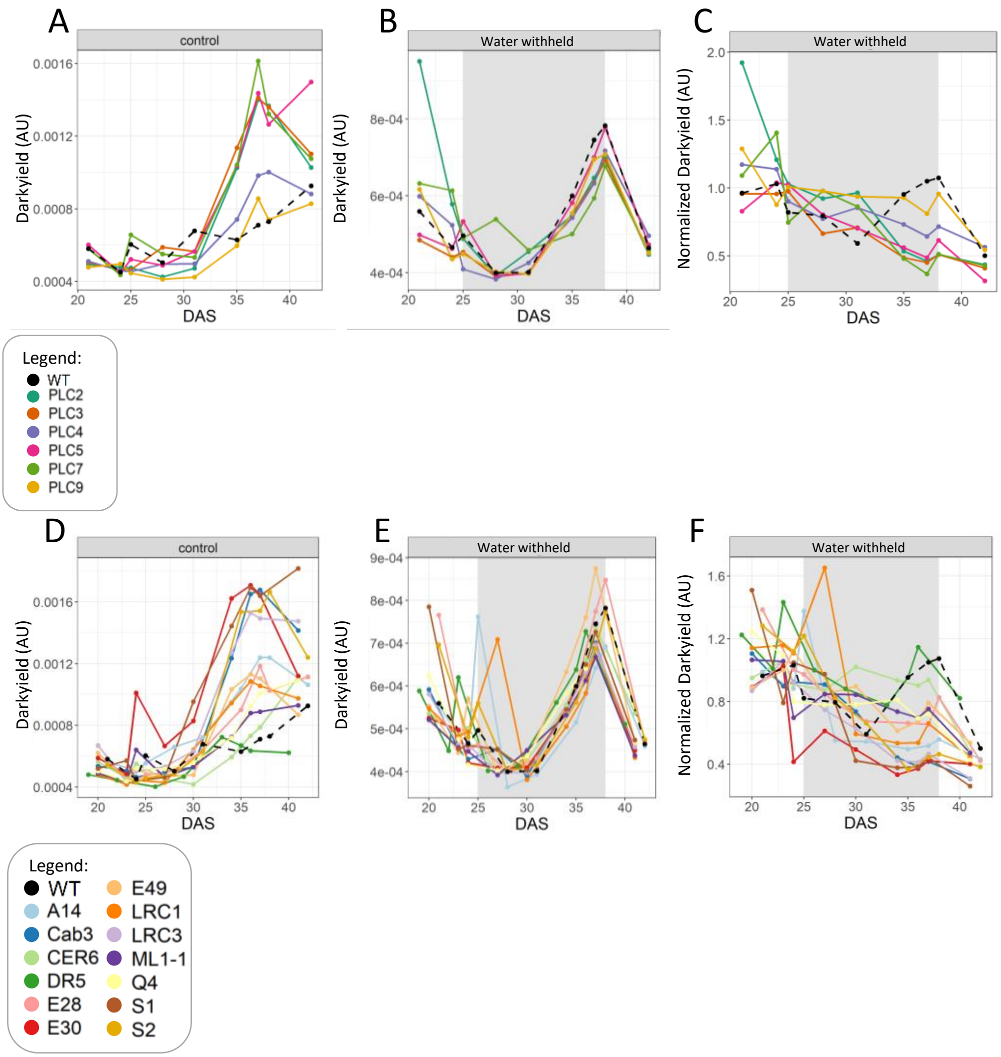


**Supplemental Figure 2.** Darkyield of plants at control- and water limiting conditions. WT plants were compared to independent transgenic lines in duplo, using 30 replicas each.

A) Darkyield of PLC-OEs at control conditions.

B) Darkyield of PLC-OEs during water withholding, grey shading indicates the period of water withholding from 25 – 38 DAS.

C) Water withholding conditions of PLC-OEs normalized to control conditions.

D) Darkyield of TSEP at control conditions.

E) Darkyield of TSEP during water withholding, grey shading indicates the period of water withholding from 25 – 38 DAS.

F) Water withholding conditions normalized to control conditions.

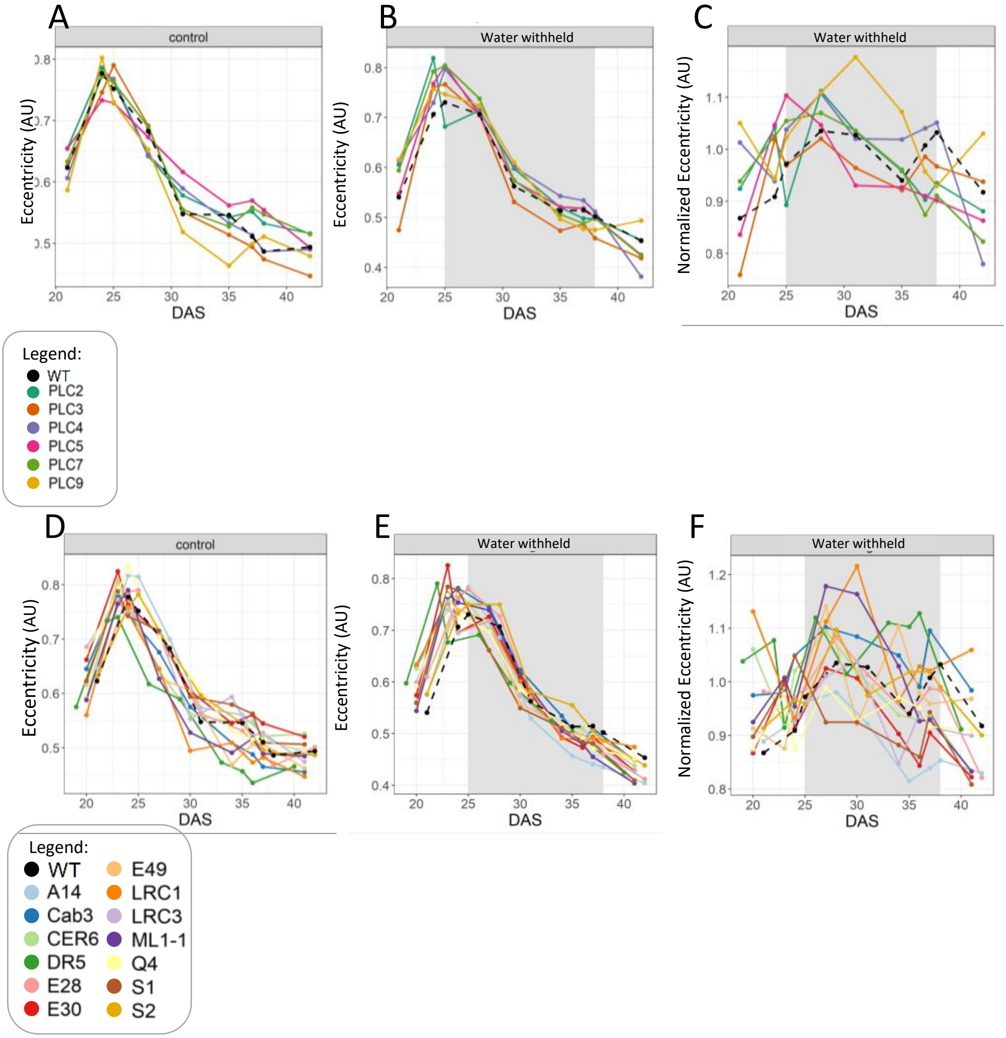


**Supplemental Figure 3.** Eccentricity of plants at control- and water limiting conditions. WT plants were compared to independent transgenic lines in duplo, using 30 replicas each.

A) Eccentricity pf PLC-OEs at control conditions.

B) Eccentricity of PLC-OEs during water withholding, grey shading indicates the period of water withholding from 25 – 38 DAS.

C) Water withholding conditions of PLC-OEs normalized to control conditions.

D) Eccentricity of TSEP at control conditions.

E) Eccentricity of TSEP during water withholding, grey shading indicates the period of water withholding from 25 – 38 DAS.

F) Water withholding conditions normalized to control conditions.

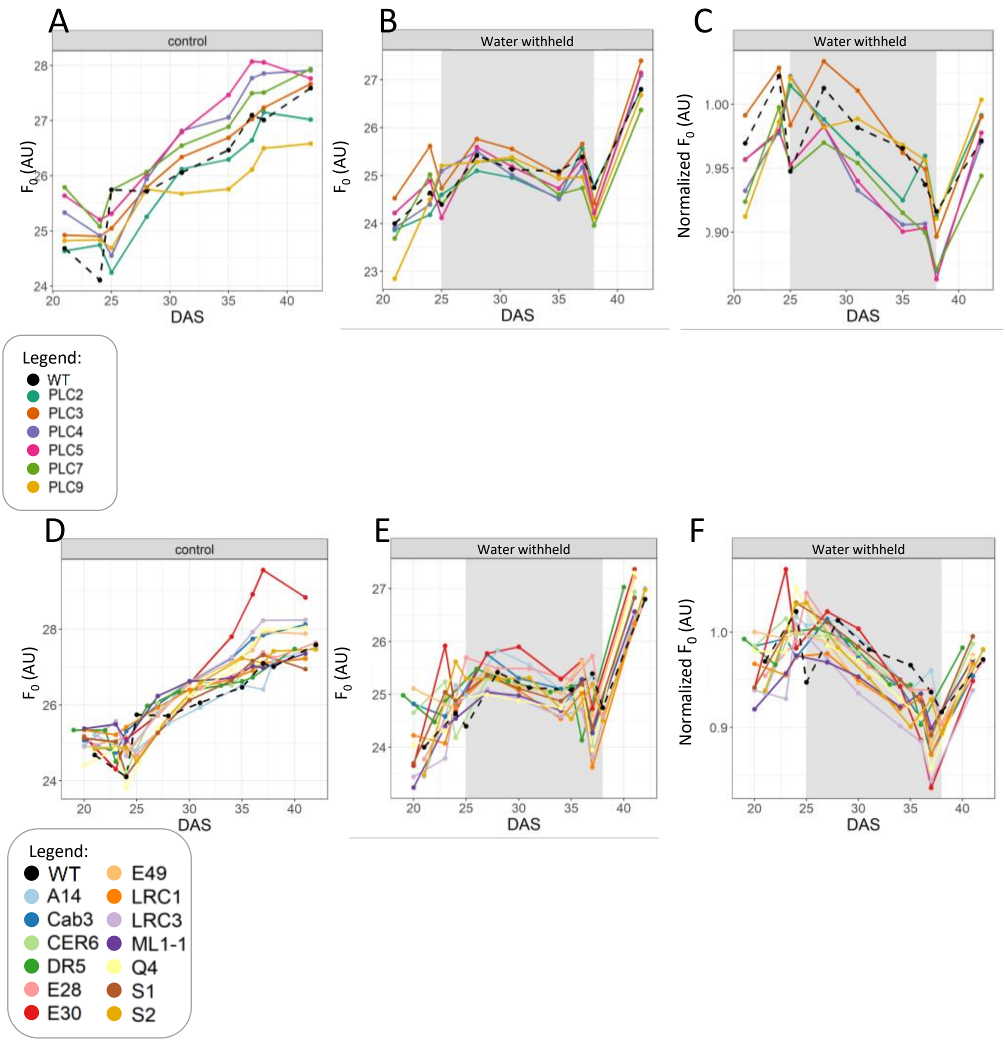


**Supplemental Figure 4.** $F_0$ of plants at control- and water limiting conditions.
WT plants were compared to independent transgenic lines in duplo, using 30 replicas each.

A) $F_0$ pf PLC-OEs at control conditions.

B) $F_0$ of PLC-OEs during water withholding, grey shading indicates the period of water withholding from 25 – 38 DAS.

C) Water withholding conditions of PLC-OEs normalized to control conditions.

D) $F_0$ of TSEP at control conditions.

E) $F_0$ of TSEP during water withholding, grey shading indicates the period of water withholding from 25 – 38 DAS.

F) Water withholding conditions normalized to control conditions.

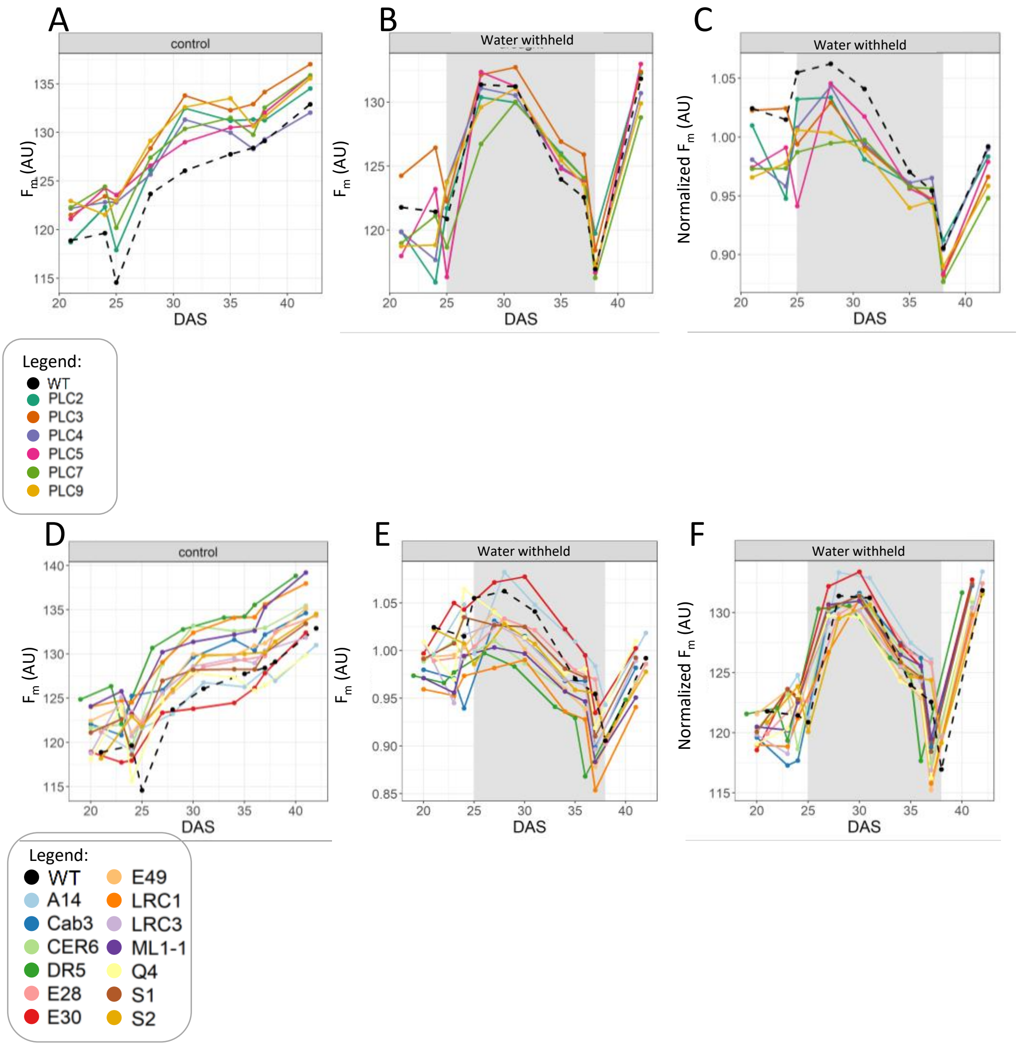


**Supplemental Figure 5.** $F_m$ of plants at control- and water limiting conditions.
WT plants were compared to independent transgenic lines in duplo, using 30 replicas each.

A) $F_m$ pf PLC-OEs at control conditions.

B) $F_m$ of PLC-OEs during water withholding, grey shading indicates the period of water withholding from 25 – 38 DAS.

C) Water withholding conditions of PLC-OEs normalized to control conditions.

D) $F_m$ of TSEP at control conditions.

E) $F_m$ of TSEP during water withholding, grey shading indicates the period of water withholding from 25 – 38 DAS.

F) Water withholding conditions normalized to control conditions.

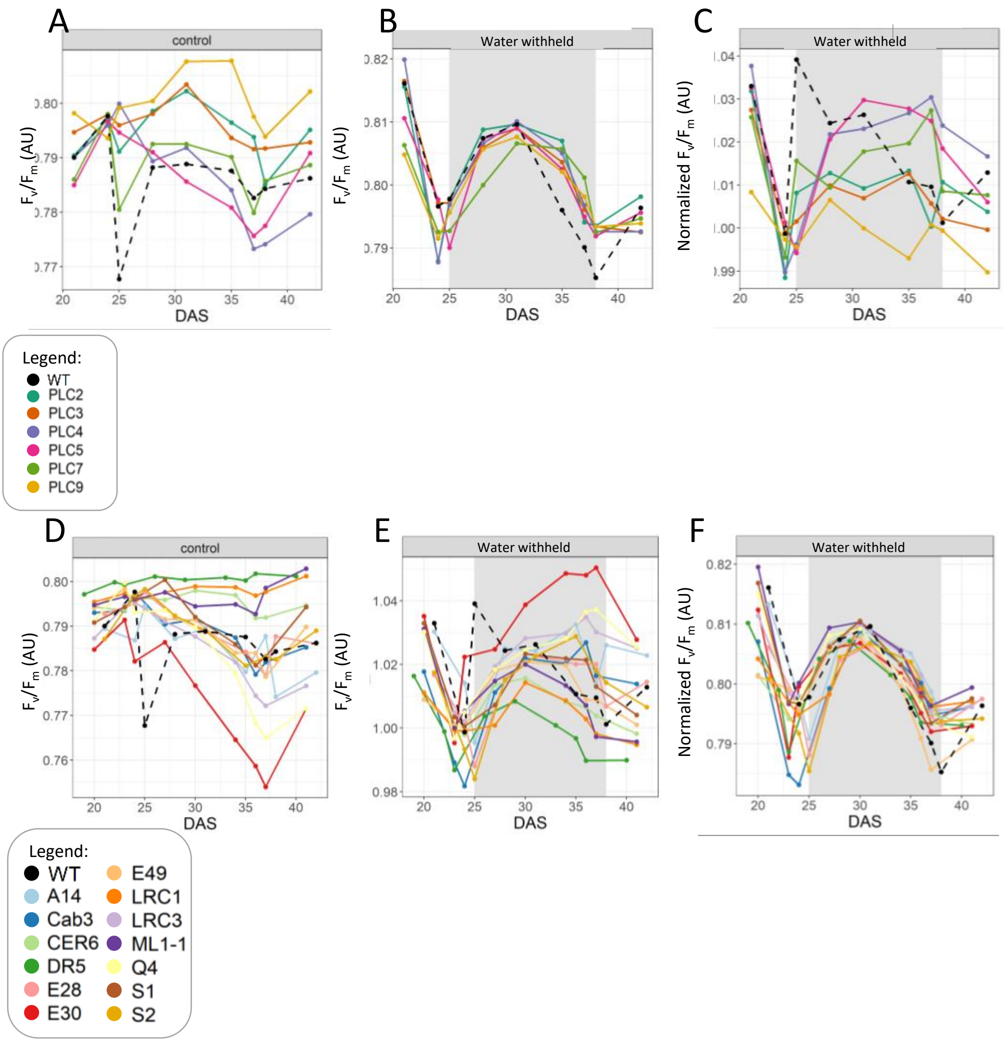


**Supplemental Figure 6.** $F_v/F_m$ ratios of plants at control- and water limiting conditions. WT plants were compared to independent transgenic lines in duplo, using 30 replicas each.

A) $F_v/F_m$ ratios of PLC-OEs at control conditions.

B) $F_v/F_m$ ratios of PLC-OEs during water withholding, grey shading indicates the period of water withholding from 25 – 38 DAS.

C) Water withholding conditions of PLC-OEs normalized to control conditions.

D) $F_v/F_m$ ratios of TSEP at control conditions.

E) $F_v/F_m$ ratios of TSEP during water withholding, grey shading indicates the period of water withholding from 25 – 38 DAS.

F) Water withholding conditions normalized to control conditions.

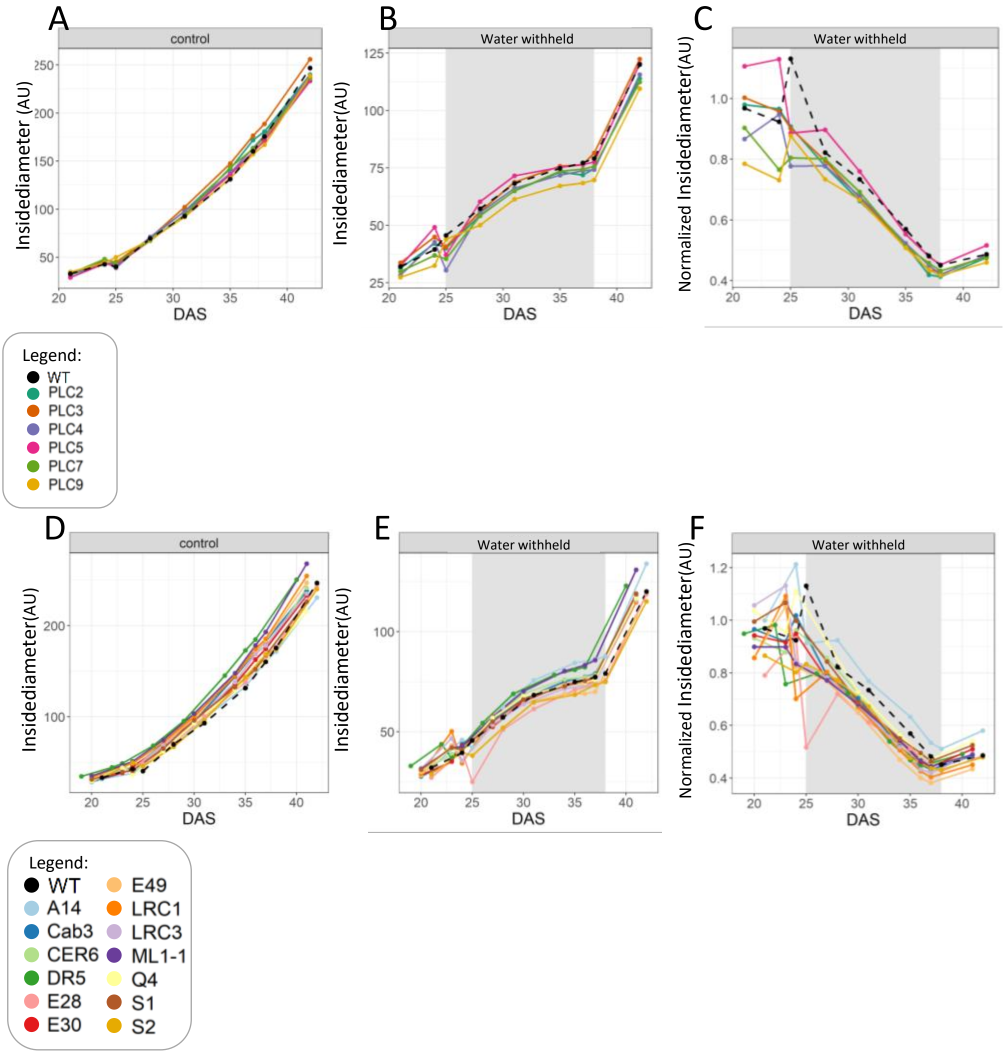


**Supplemental Figure 7.** Insidediameter of plants at control- and water limiting conditions.

WT plants were compared to independent transgenic lines in duplo, using 30 replicas each.

A) Insidediameter of PLC-OEs at control conditions.

B) Insidediameter of PLC-OEs during water withholding, grey shading indicates the period of water withholding from 25 – 38 DAS.

C) Water withholding conditions of PLC-OEs normalized to control conditions.

D) Insidediameter of TSEP at control conditions.

E) Insidediameter of TSEP during water withholding, grey shading indicates the period of water withholding from 25 – 38 DAS.

F) Water withholding conditions normalized to control conditions.

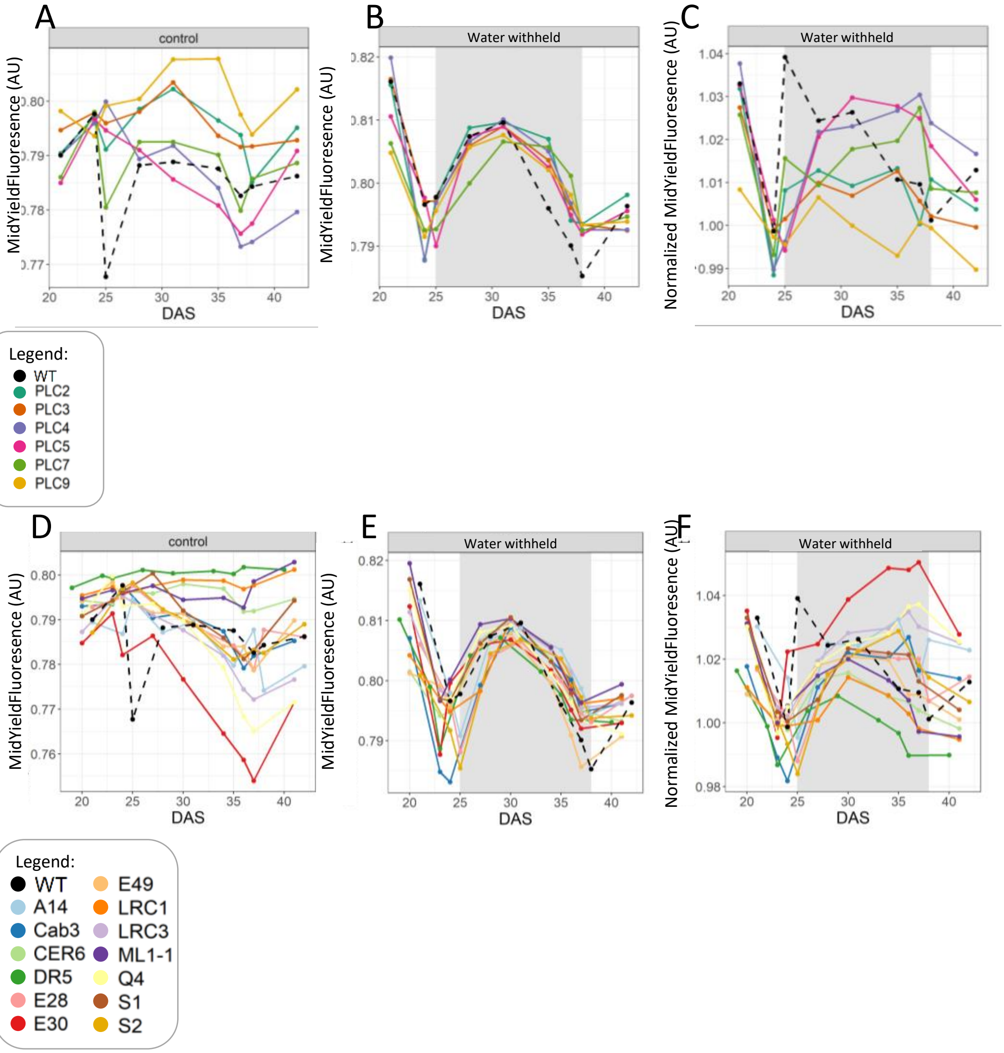


**Supplemental Figure 8.** midyieldfluoresence of plants at control- and water limiting conditions.

WT plants were compared to independent transgenic lines in duplo, using 30 replicas each.

A) midyieldfluoresence of PLC-OEs at control conditions.

B) midyieldfluoresence of PLC-OEs during water withholding, grey shading indicates the period of water withholding from 25 – 38 DAS.

C) Water withholding conditions of PLC-OEs normalized to control conditions.

D) midyieldfluoresence of TSEP at control conditions.

E) midyieldfluoresence of TSEP during water withholding, grey shading indicates the period of water withholding from 25 – 38 DAS.

F) Water withholding conditions normalized to control conditions.

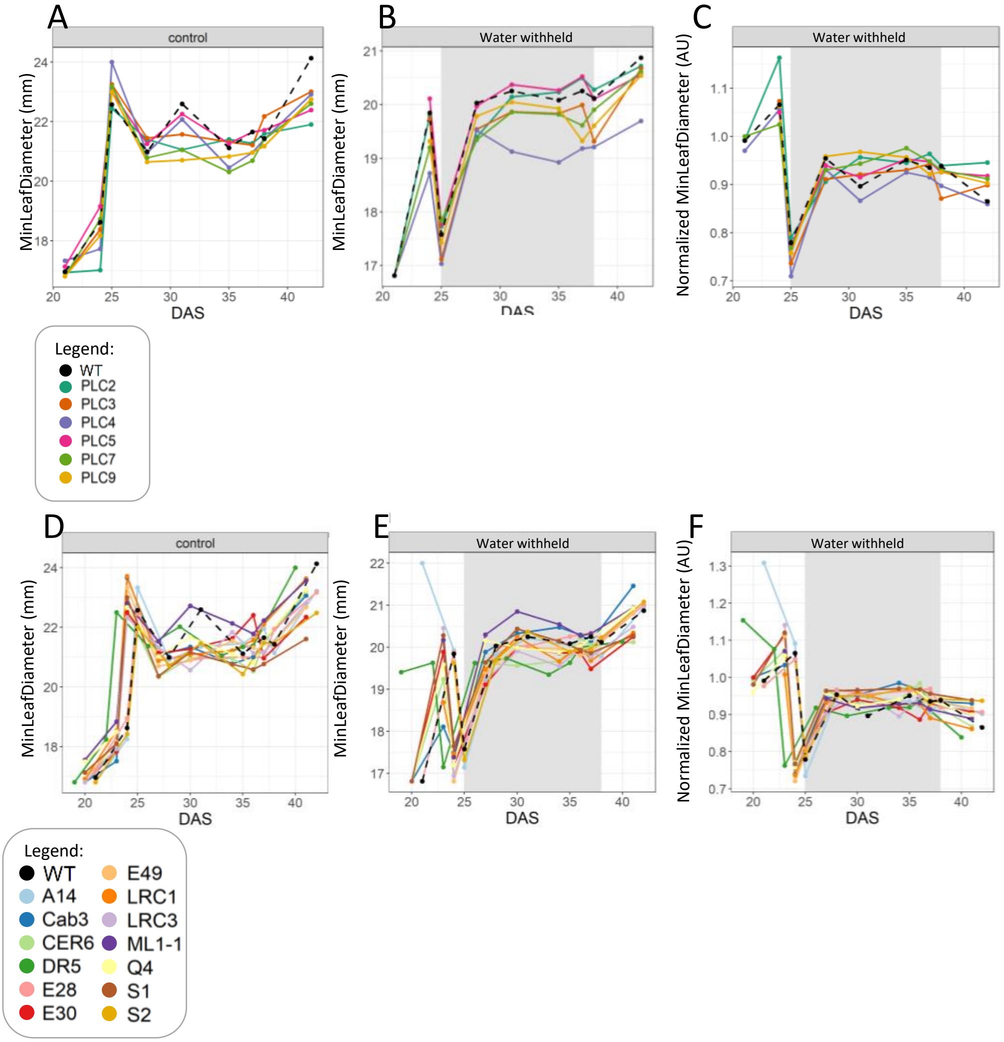


**Supplemental Figure 9.** Minimal leaf diameter of plants at control- and water limiting conditions.

WT plants were compared to independent transgenic lines in duplo, using 30 replicas each.

A) Minimal leaf diameter of PLC-OEs at control conditions.

B) Minimal leaf diameter of PLC-OEs during water withholding, grey shading indicates the period of water withholding from 25 – 38 DAS.

C) Water withholding conditions of PLC-OEs normalized to control conditions.

D) Minimal leaf diameter of TSEP at control conditions.

E) Minimal leaf diameter of TSEP during water withholding, grey shading indicates the period of water withholding from 25 – 38 DAS.

F) Water withholding conditions normalized to control conditions.

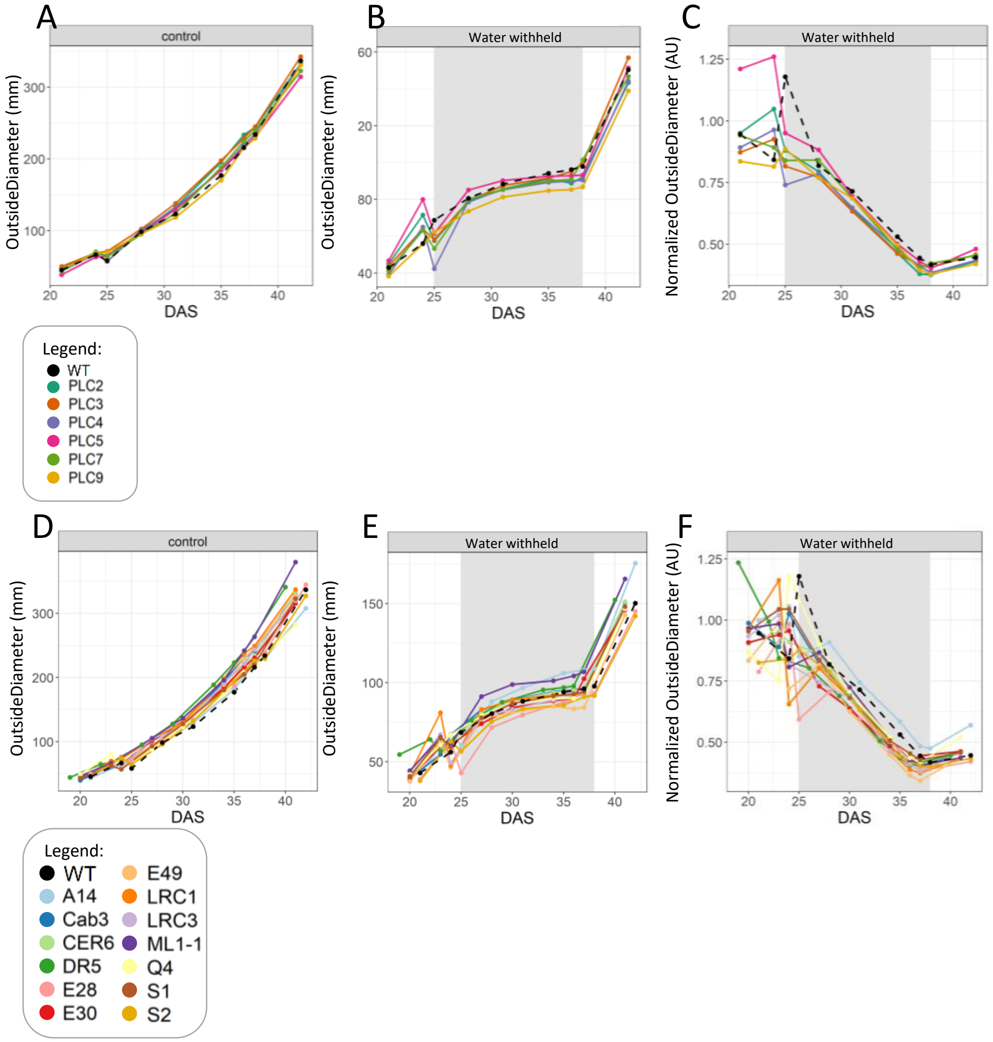


**Supplemental Figure 10.** Outside leaf diameter of plants at control- and water limiting conditions.

WT plants were compared to independent transgenic lines in duplo, using 30 replicas each.

A) Outside leaf diameter of PLC-OEs at control conditions.

B) Outside leaf diameter of PLC-OEs during water withholding, grey shading indicates the period of water withholding from 25 – 38 DAS.

C) Water withholding conditions of PLC-OEs normalized to control conditions.

D) Outside leaf diameter of TSEP at control conditions.

E) Outside leaf diameter of TSEP during water withholding, grey shading indicates the period of water withholding from 25 – 38 DAS.

F) Water withholding conditions normalized to control conditions.

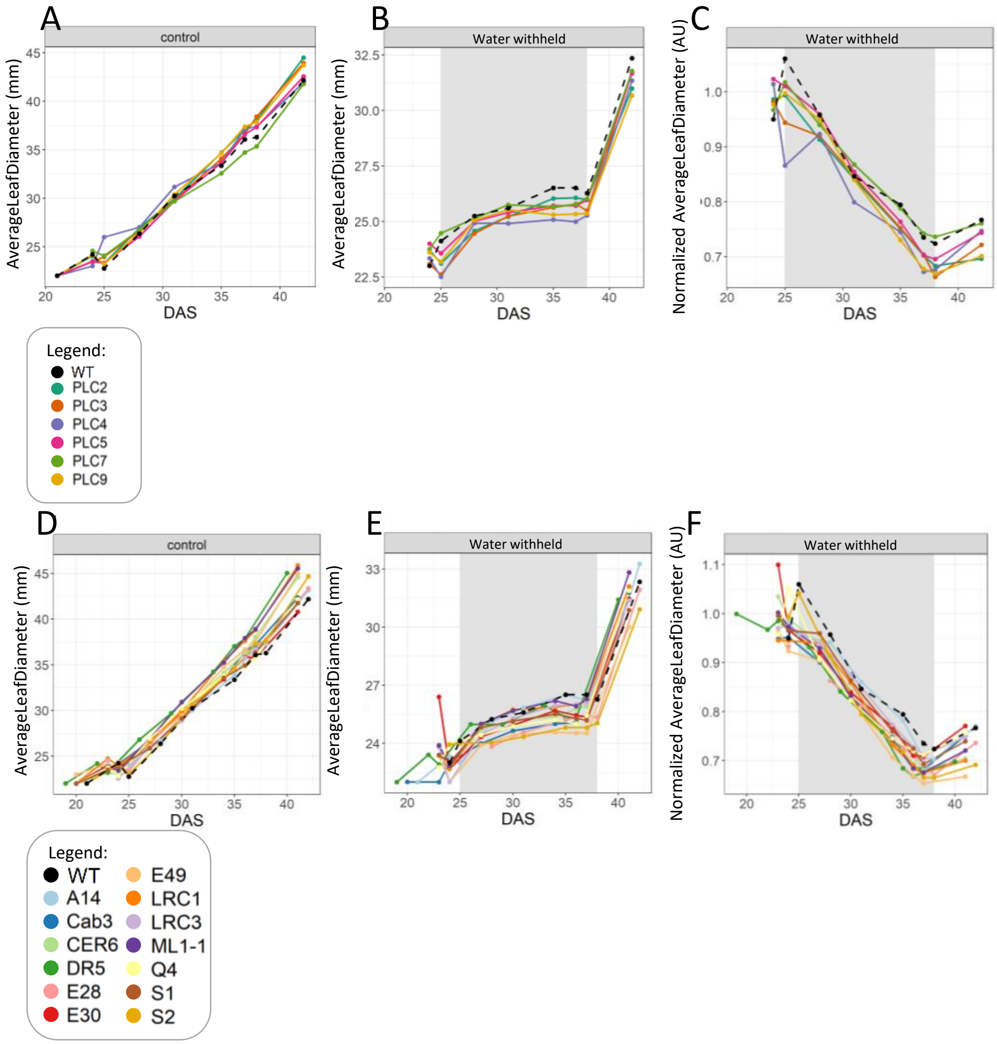


**Supplemental Figure 11.** Average leaf diameter of plants at control- and water limiting conditions.
WT plants were compared to independent transgenic lines in duplo, using 30 replicas each.
A) Average leaf diameter of PLC-OEs at control conditions.
B) Average leaf diameter of PLC-OEs during water withholding, grey shading indicates the period of water withholding from 25 – 38 DAS.
C) Water withholding conditions of PLC-OEs normalized to control conditions.
D) Average leaf diameter of TSEP at control conditions.
E) Average leaf diameter of TSEP during water withholding, grey shading indicates the period of water withholding from 25 – 38 DAS.
F) Water withholding conditions normalized to control conditions.

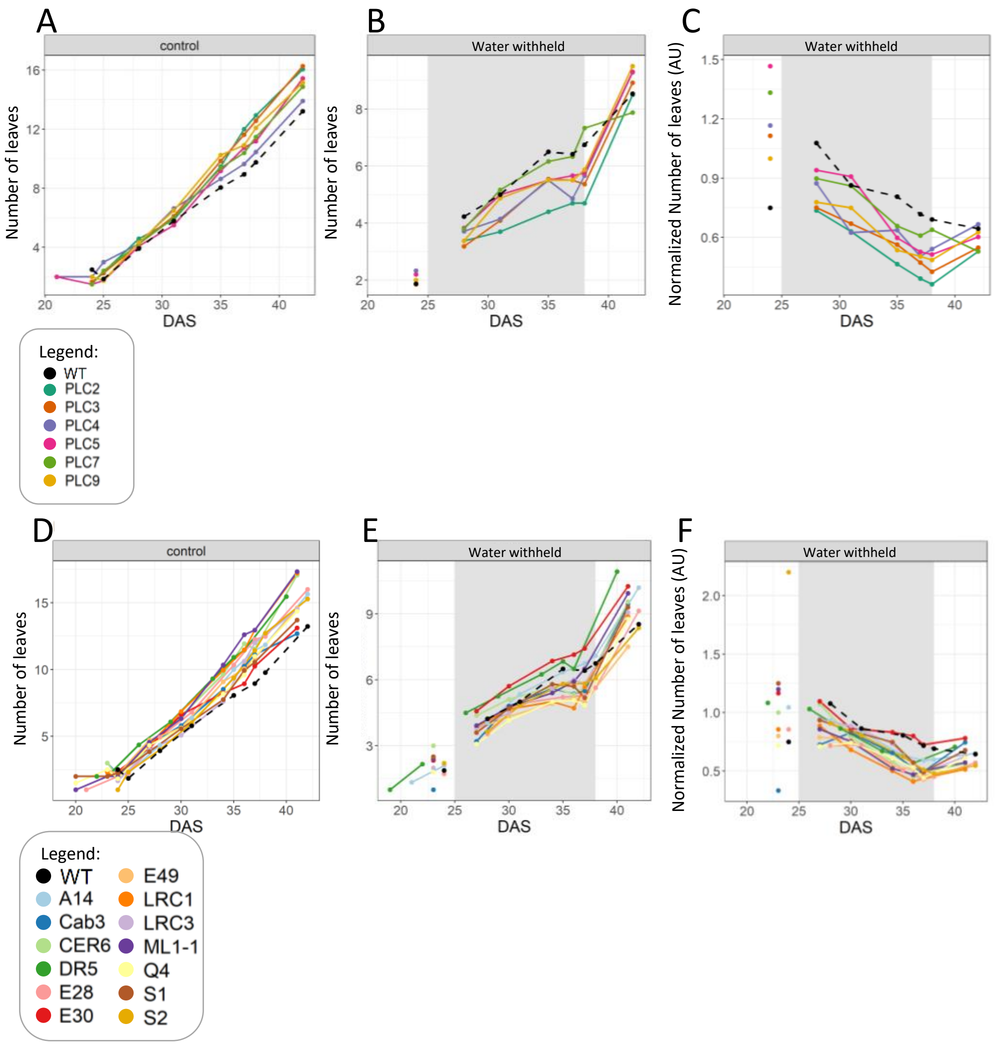


**Supplemental Figure 12.** Number of leaves of plants at control- and water limiting conditions.
WT plants were compared to independent transgenic lines in duplo, using 30 replicas each.
A) Number of leaves of PLC-OEs at control conditions.
B) Number of leaves of PLC-OEs during water withholding, grey shading indicates the period of water withholding from 25 – 38 DAS.
C) Water withholding conditions of PLC-OEs normalized to control conditions.
D) Number of leaves of TSEP at control conditions.
E) Number of leaves of TSEP during water withholding, grey shading indicates the period of water withholding from 25 – 38 DAS.
F) Water withholding conditions normalized to control conditions.

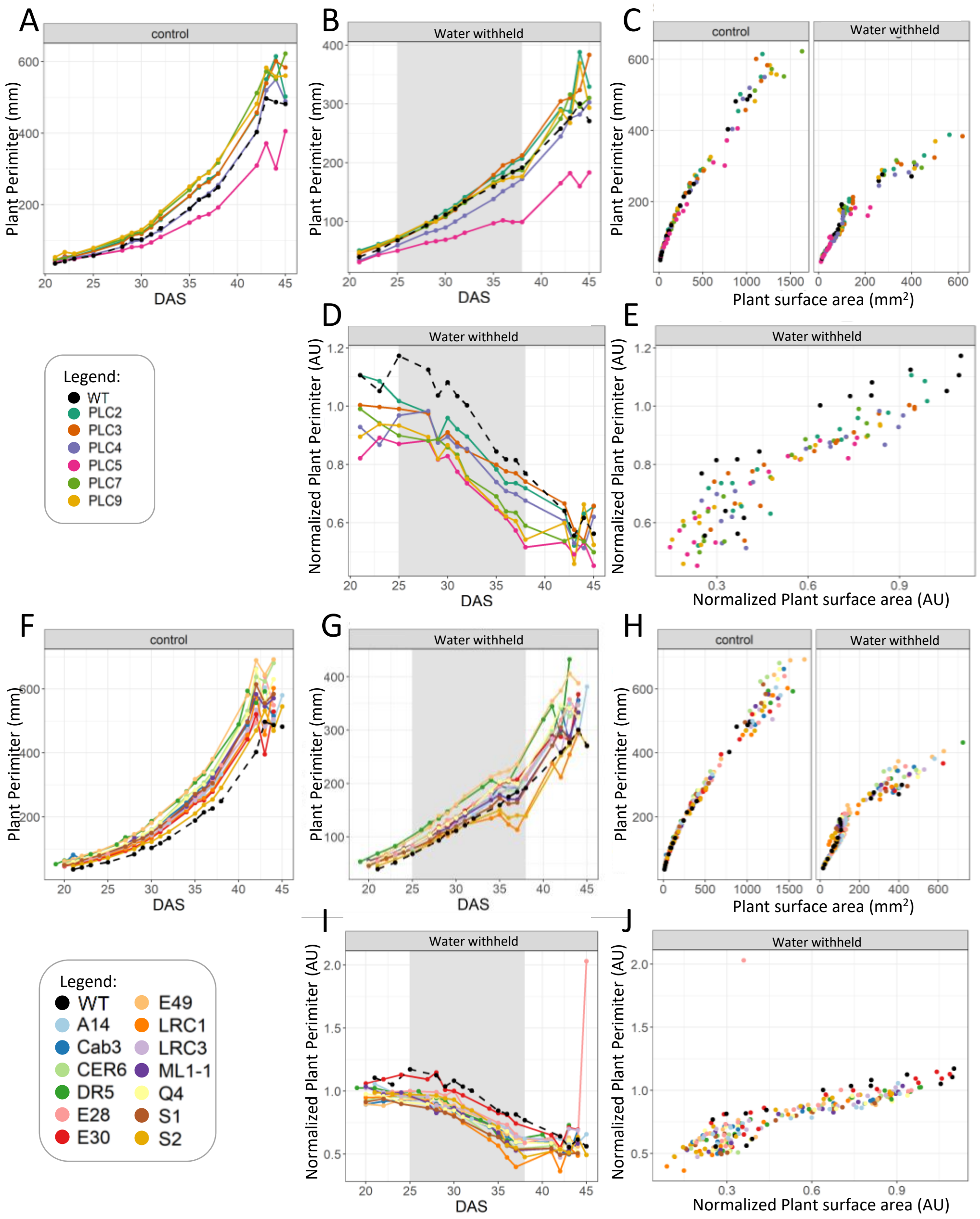


**Supplemental figure 13.** Plant perimeter at control- and water limiting conditions.
The Plant perimeter was measured for:
A, F) PLC-OE and TSEP grown at control conditions.
B, G) PLC-OE and TSEP at water-withholding conditions, with grey shading indicating period of water withholding (25-38 DAS).
C, H) PLC-OE and TSEP in Water-withholding conditions normalized to control per day
D, I ) PLC-OE and TSEP in Water withholding conditions normalized to control conditions.
E, J) PLC-OE and TSEP in water withholding conditions normalized to Projected plant surface area

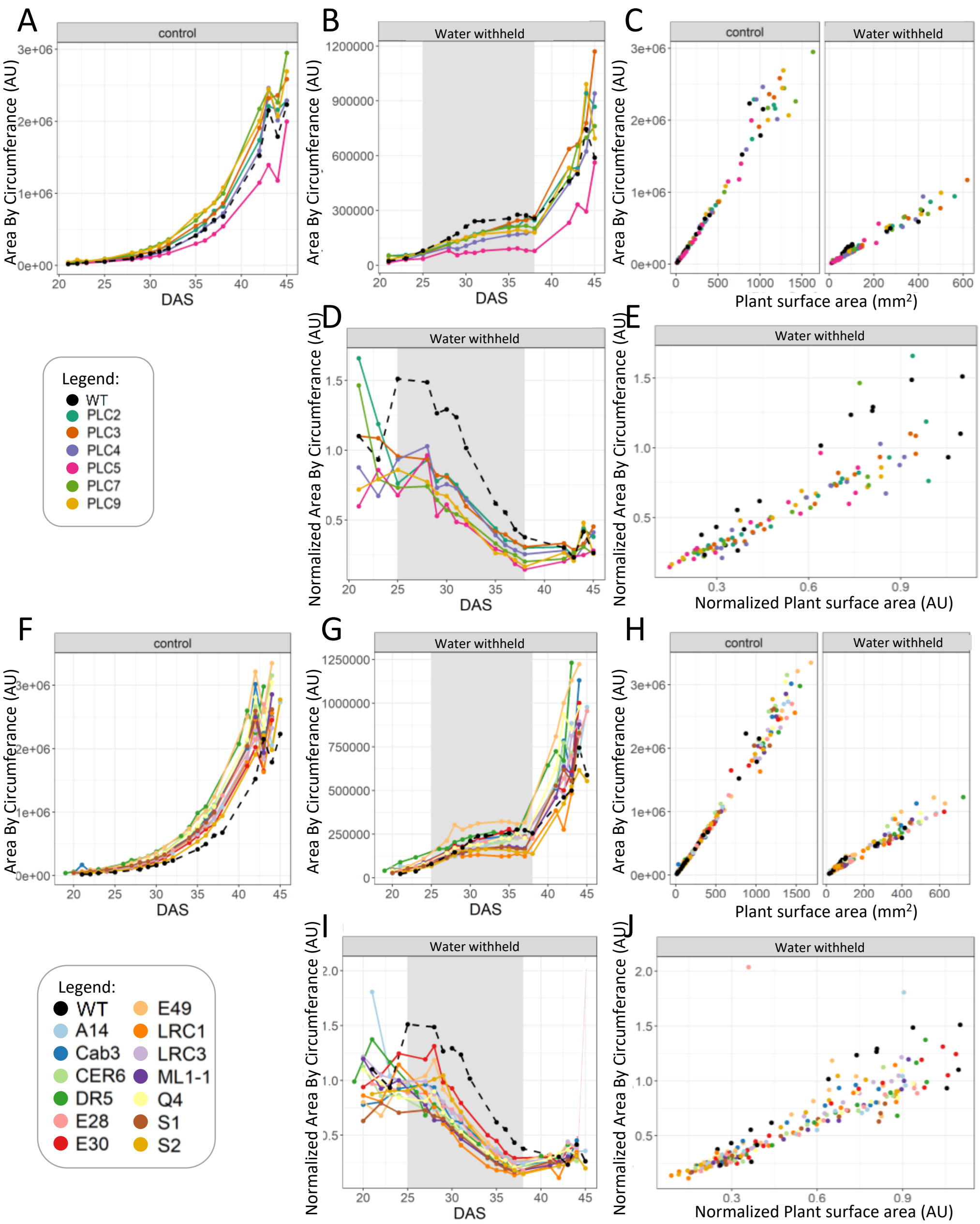


**Supplemental figure 14.** Area by circumference at control- and water limiting conditions. The Area by circumference was measured for:

A, F) PLC-OE and TSEP grown at control conditions.

B, G) PLC-OE and TSEP at water-withholding conditions, with grey shading indicating period of water withholding (25-38 DAS).

C, H) PLC-OE and TSEP in Water-withholding conditions normalized to control per day

D, I ) PLC-OE and TSEP in Water withholding conditions normalized to control conditions.

E, J) PLC-OE and TSEP in water withholding conditions normalized to Projected plant surface area

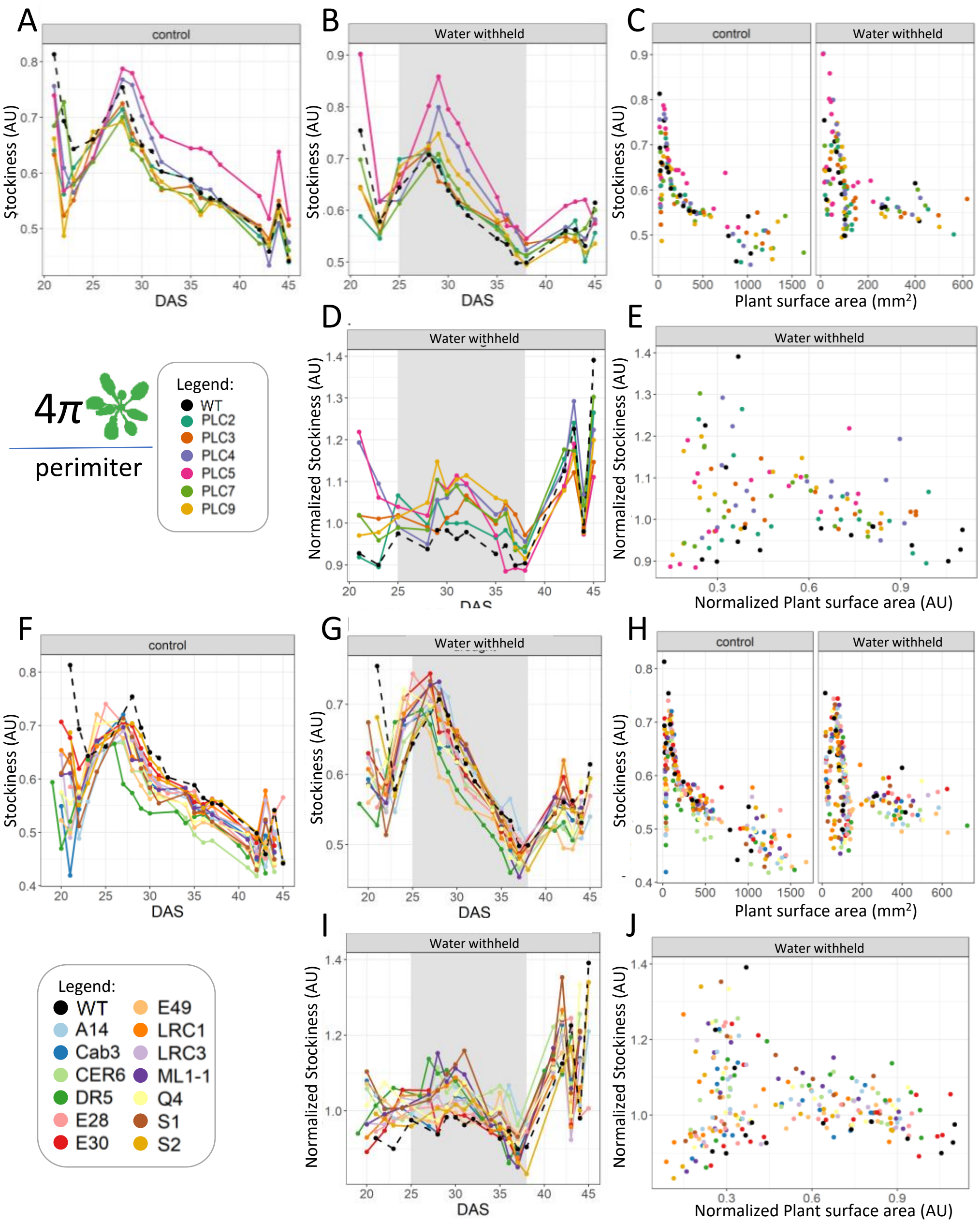


**Supplemental figure 15.** Stockiness at control- and water limiting conditions.
The stockiness was measured for:
A, F) PLC-OE and TSEP grown at control conditions.
B, G) PLC-OE and TSEP at water-withholding conditions, with grey shading indicating period of water withholding (25-38 DAS).
C, H) PLC-OE and TSEP in Water-withholding conditions normalized to control per day
D, I ) PLC-OE and TSEP in Water withholding conditions normalized to control conditions.
E, J) PLC-OE and TSEP in water withholding conditions normalized to Projected plant surface area

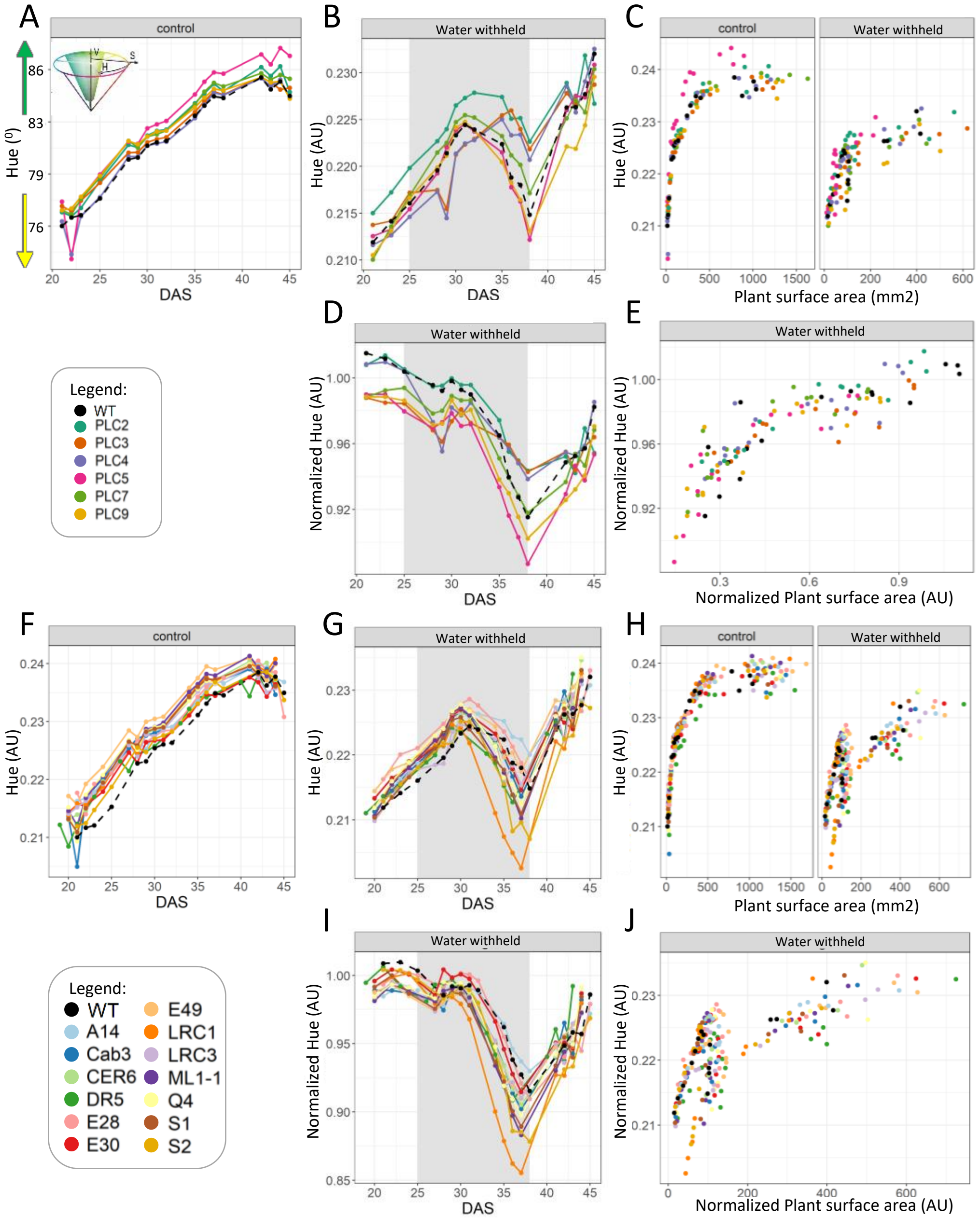


**Supplemental figure 16.** Hue at control- and water limiting conditions.
The Hue was measured for :
A, F) PLC-OE and TSEP grown at control conditions.
B, G) PLC-OE and TSEP at water-withholding conditions, with grey shading indicating period of water withholding (25-38 DAS).
C, H) PLC-OE and TSEP in Water-withholding conditions normalized to control per day
D, I ) PLC-OE and TSEP in Water withholding conditions normalized to control conditions.
E, J) PLC-OE and TSEP in water withholding conditions normalized to Projected plant surface area

### Brightness PLC

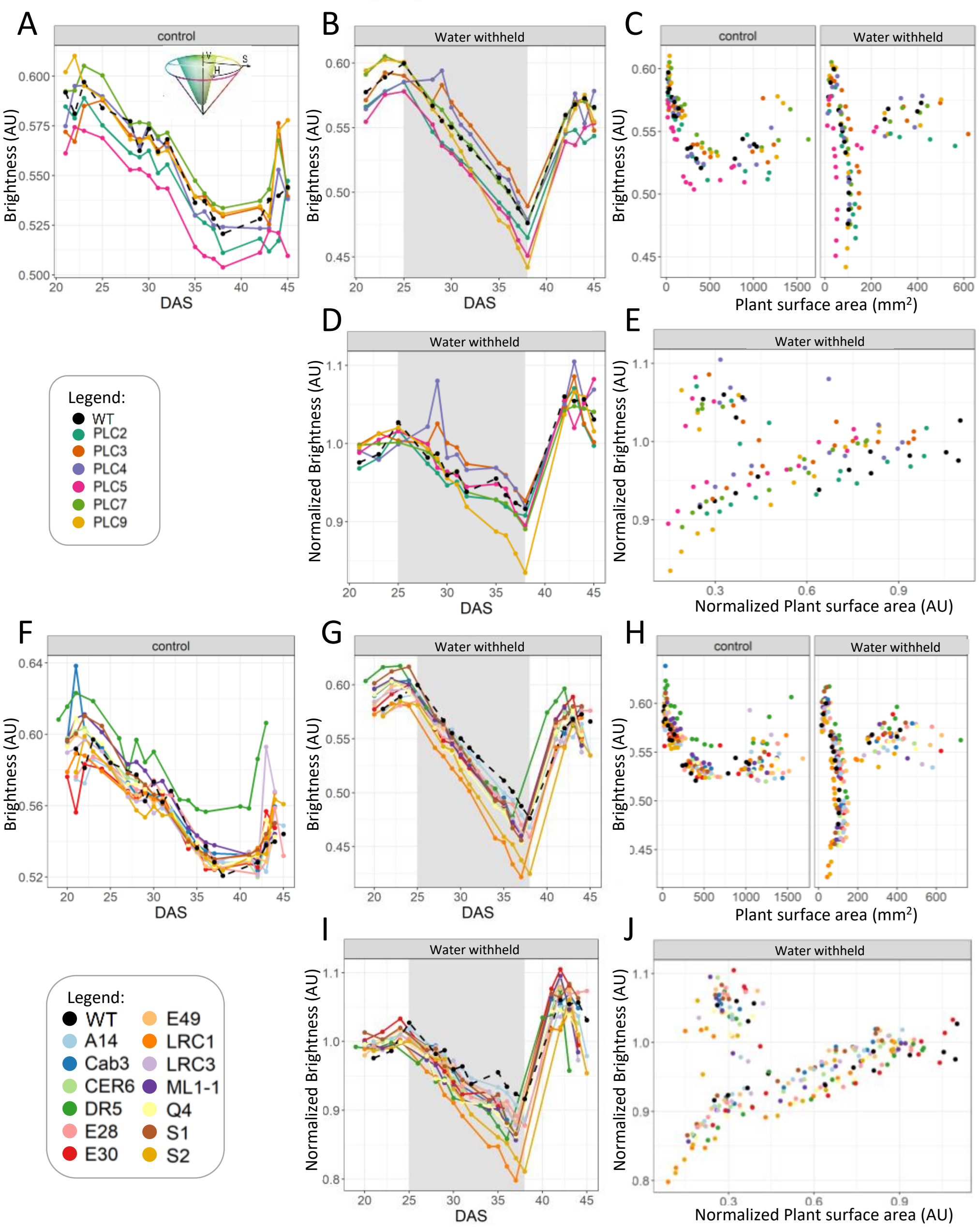


**Supplemental figure 17.** brightness at control- and water limiting conditions.
The brightness was measured for :
A, F) PLC-OE and TSEP grown at control conditions.
B, G) PLC-OE and TSEP at water-withholding conditions, with grey shading indicating period of water withholding (25-38 DAS).
C, H) PLC-OE and TSEP in Water-withholding conditions normalized to control per day
D, I ) PLC-OE and TSEP in Water withholding conditions normalized to control conditions.
E, J) PLC-OE and TSEP in water withholding conditions normalized to Projected plant surface area

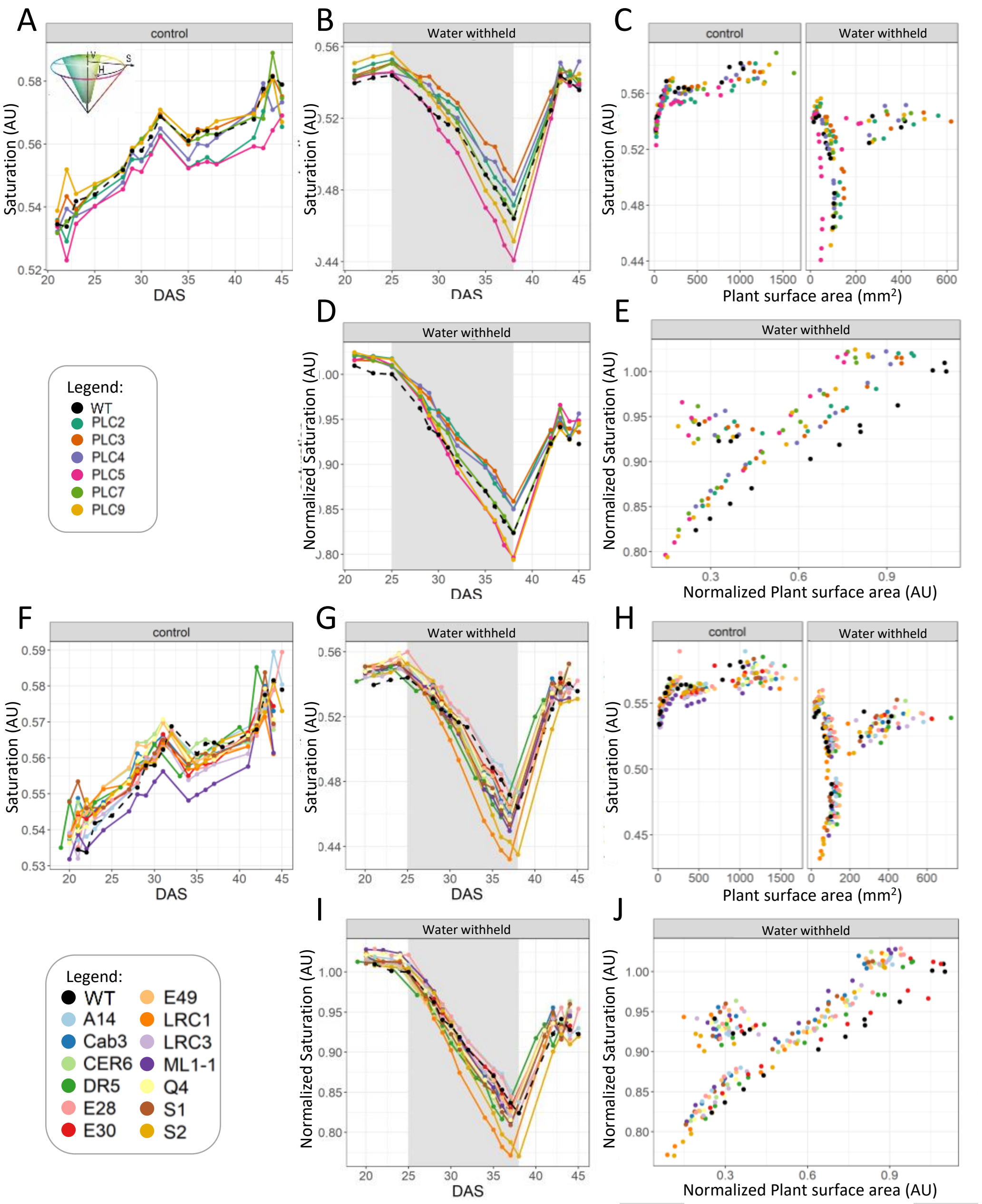


**Supplemental figure 18.** Saturation at control- and water limiting conditions.
The Saturation was measured for :
A, F) PLC-OE and TSEP grown at control conditions.
B, G) PLC-OE and TSEP at water-withholding conditions, with grey shading indicating period of water withholding (25-38 DAS).
C, H) PLC-OE and TSEP in Water-withholding conditions normalized to control per day
D, I ) PLC-OE and TSEP in Water withholding conditions normalized to control conditions.
E, J) PLC-OE and TSEP in water withholding conditions normalized to Projected plant surface area

### Figure 17

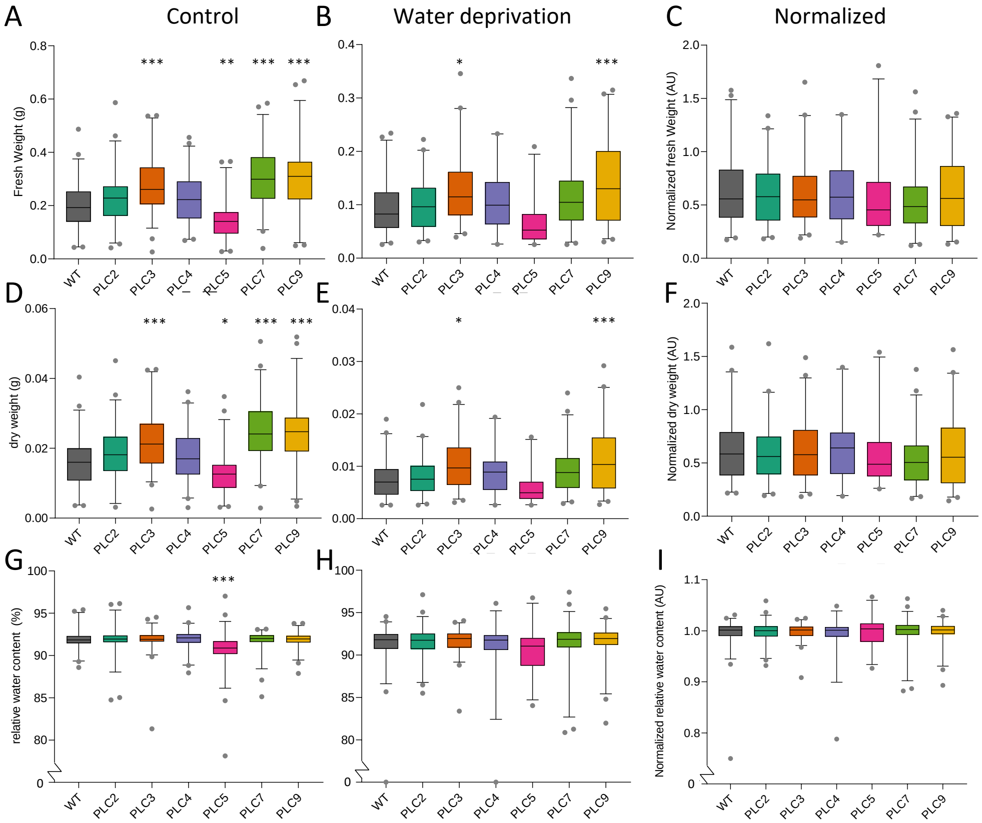


**Supplemental figure 19**. Fresh- and dry weight and relative water content in WT and PLC-OE lines.
Plants were harvested and measured at 45 DAS. Water-stressed plants were deprived of water for 13 days, and left to recover for seven days before harvest. After measuring fresh weight (FW), plants were dried for 48 hours in a 70° oven and measured for their dry weight (DW). RWC was determined by the formula: (FW-DW)/FW*100%.
A-C) Fresh weights (control-, drought-, or drought normalized to control conditions, respectively).
D-F) Dry weights
G-I) RWC (A-F)
Significance was determined by one-way ANOVA, with multiple comparison test Dunnet. *P < 0.05, P**<0.01,***P < 0.001

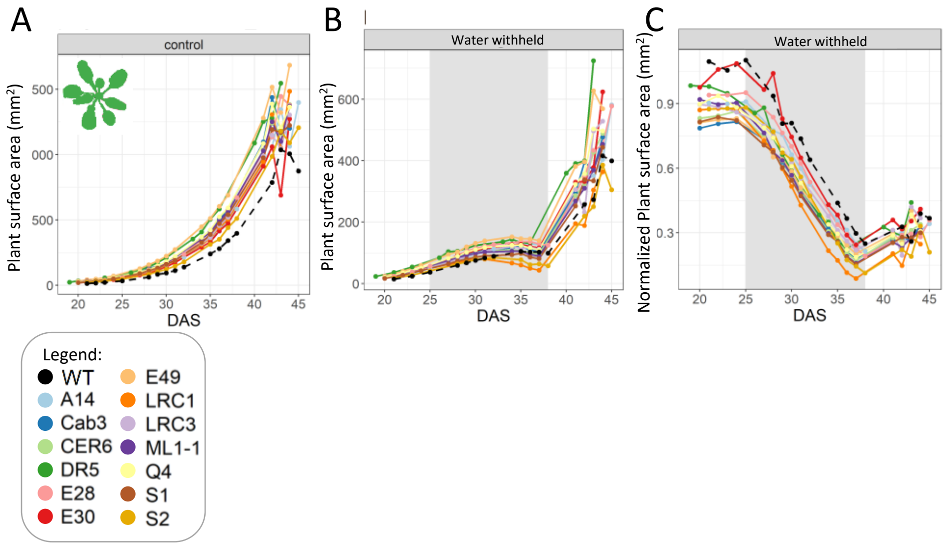


**Supplemental figure 20**. Projected plant surface area of TSEP lines under control and water limiting conditions.

Arabidopsis WT and TSEP lines plants were monitored using the GROWSCREEN-FLUORO system for indicated times. For each genotype, two different insertion lines with 30 replicates were used.

A) Projected leaf area (schematically drawn in upper left corner) at control conditions.

B) Projected leaf area at water-withholding conditions (25-37 DAS).

C) Projected leaf area at water-withholding conditions normalized to control.

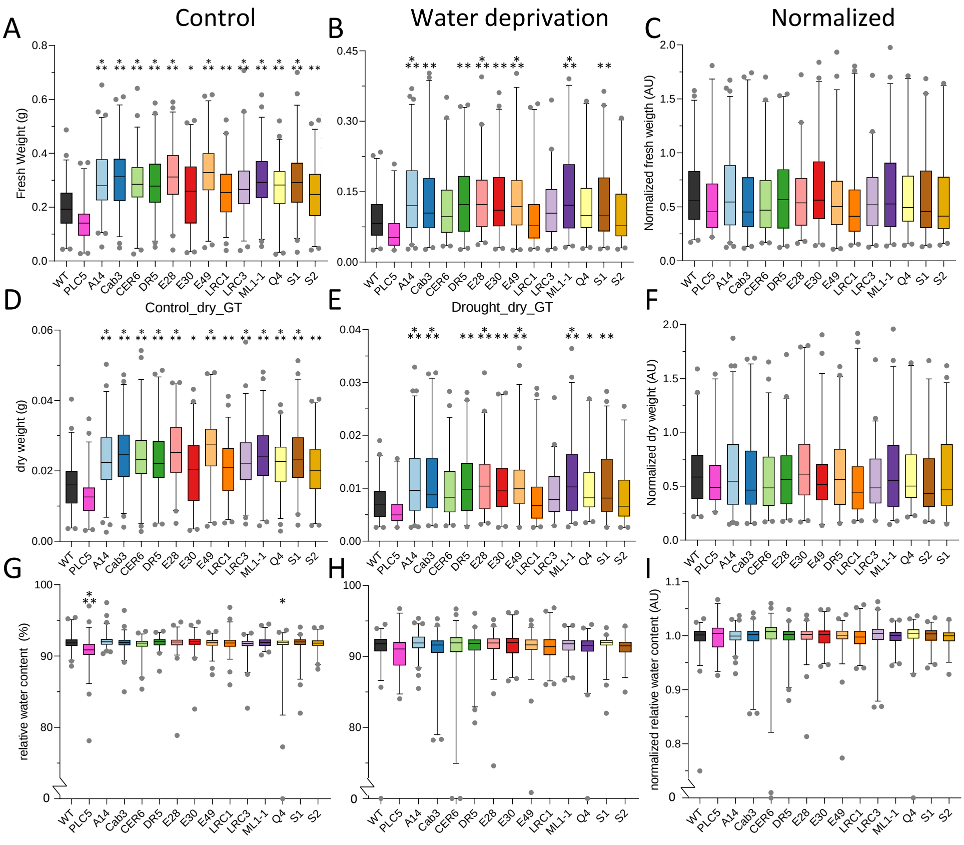


**Supplemental figure 21**. Fresh- and dry weight and relative water content in WT and TSEP lines.

Plants were harvested and measured at 45 DAS. Water-stressed plants were deprived of water for 13 days, and left to recover for seven days before harvest. After measuring fresh weight (FW), plants were dried for 48 hours in a 70° oven and measured for their dry weight (DW). RWC was determined by the formula: (FW-DW)/FW*100%.

A-C) Fresh weights (control-, drought-, or drought normalized to control conditions, respectively).

D-F) Dry weights

G-I) RWC (A-F)

Significance was determined by one-way ANOVA, with multiple comparison test Dunnet. *P < 0.05, P**<0.01,***P < 0.001

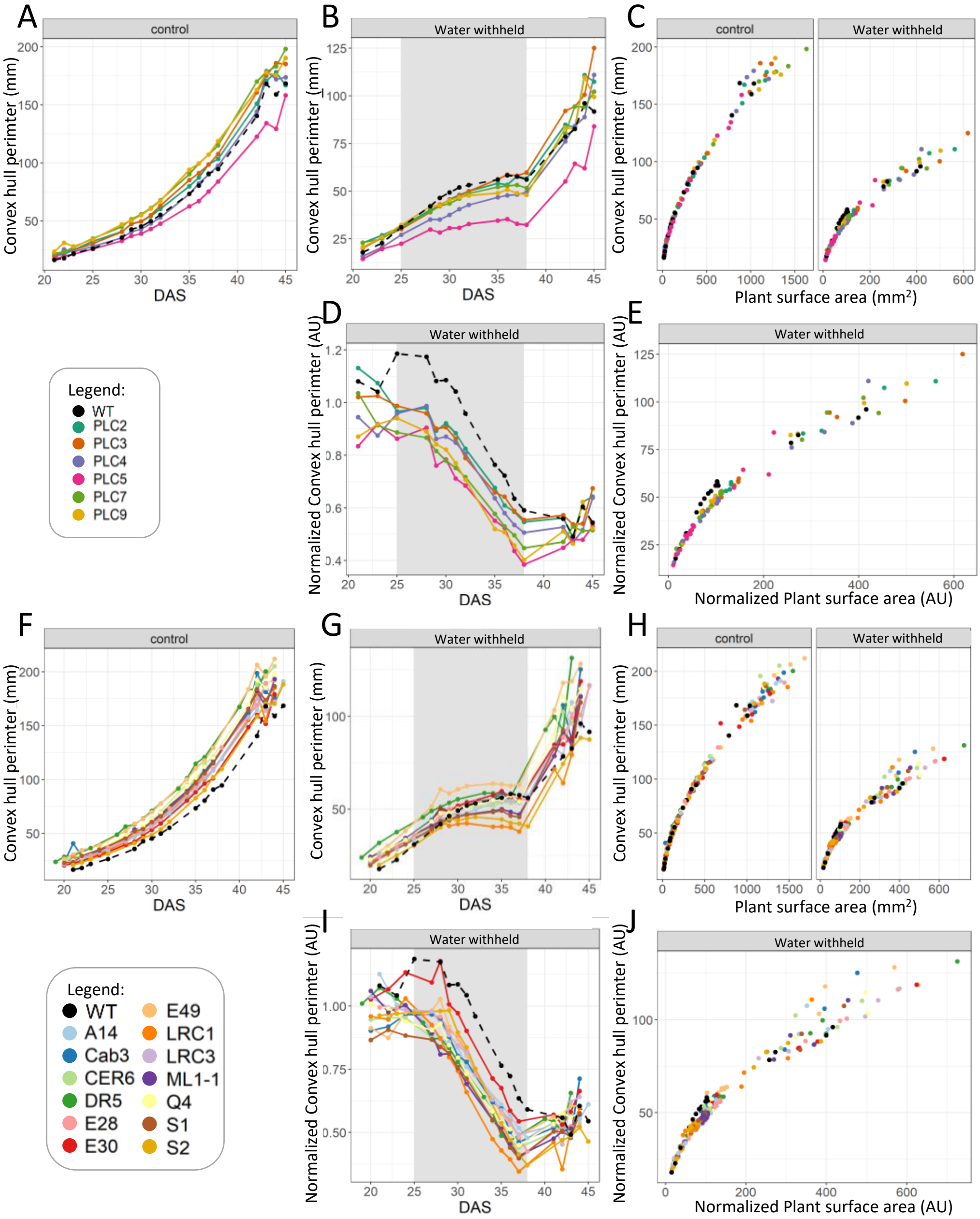


**Supplemental figure 22.** Convex hull permiter at control- and water limiting conditions. The Convex hull permiter was measured for:

A, F) PLC-OE and TSEP grown at control conditions.

B, G) PLC-OE and TSEP at water-withholding conditions, with grey shading indicating period of water withholding (25-38 DAS).

C, H) PLC-OE and TSEP in Water-withholding conditions normalized to control per day

D, I ) PLC-OE and TSEP in Water withholding conditions normalized to control conditions.

E, J) PLC-OE and TSEP in water withholding conditions normalized to Projected plant surface area

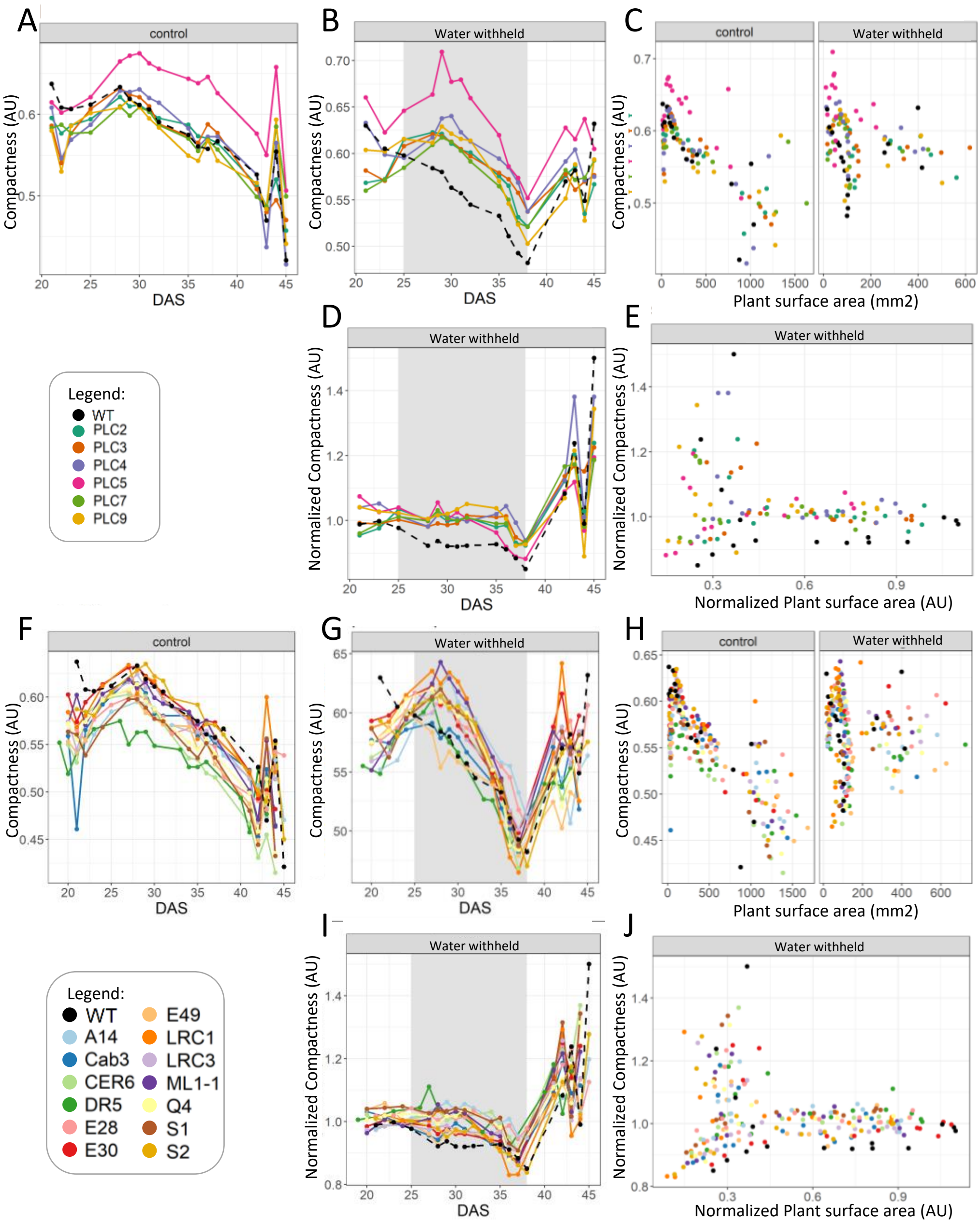


**Supplemental figure 23.** compactness at control- and water limiting conditions.
The compactness was measured for:
A, F) PLC-OE and TSEP grown at control conditions.
B, G) PLC-OE and TSEP at water-withholding conditions, with grey shading indicating period of water withholding (25-38 DAS).
C, H) PLC-OE and TSEP in Water-withholding conditions normalized to control per day
D, I ) PLC-OE and TSEP in Water withholding conditions normalized to control conditions.
E, J) PLC-OE and TSEP in water withholding conditions normalized to Projected plant surface area

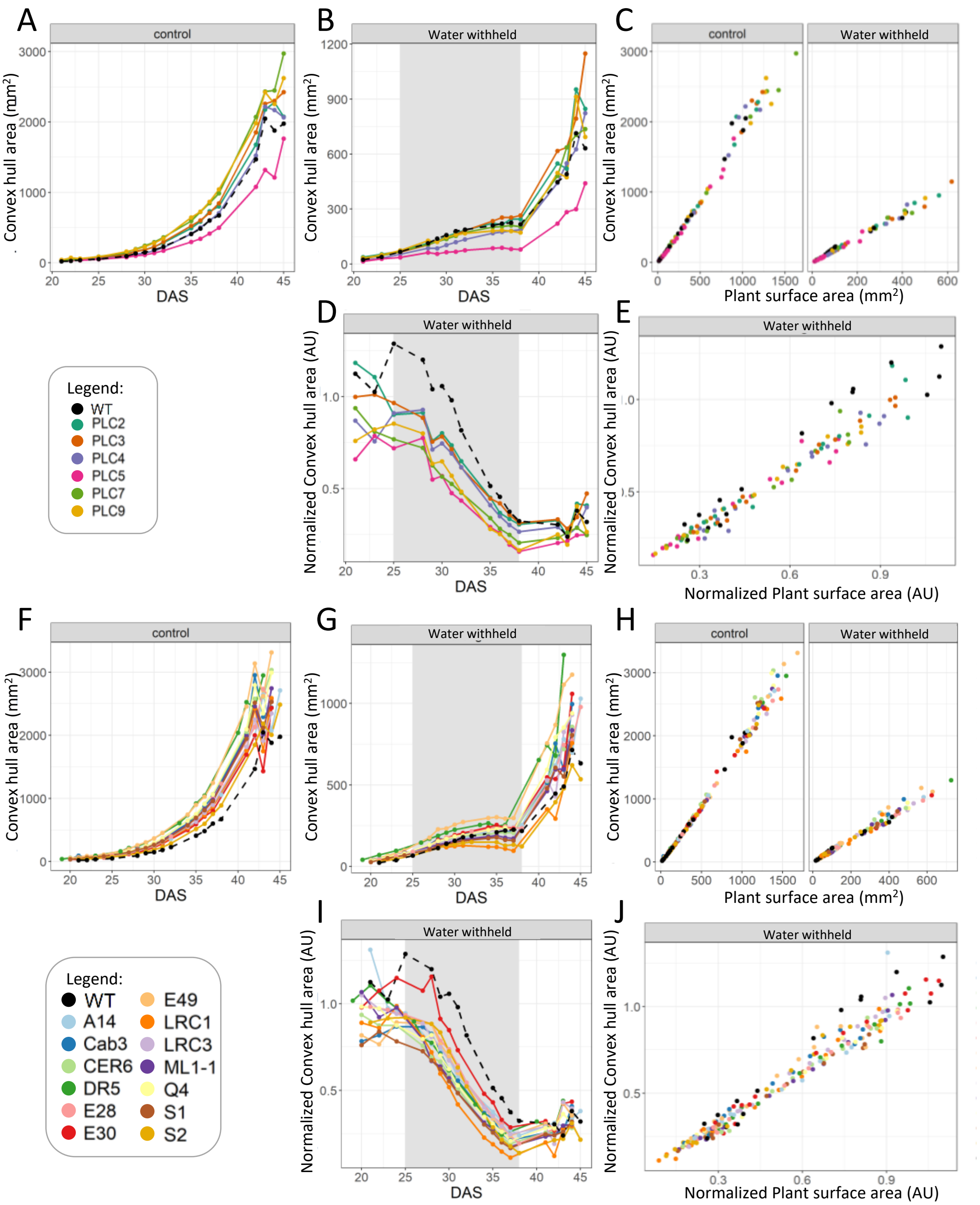


**Supplemental figure 24.** Convex hull area at control- and water limiting conditions.
The Convex hull area was measured for:
A, F) PLC-OE and TSEP grown at control conditions.
B, G) PLC-OE and TSEP at water-withholding conditions, with grey shading indicating period of water withholding (25-38 DAS).
C, H) PLC-OE and TSEP in Water-withholding conditions normalized to control per day
D, I ) PLC-OE and TSEP in Water withholding conditions normalized to control conditions.
E, J) PLC-OE and TSEP in water withholding conditions normalized to Projected plant surface area

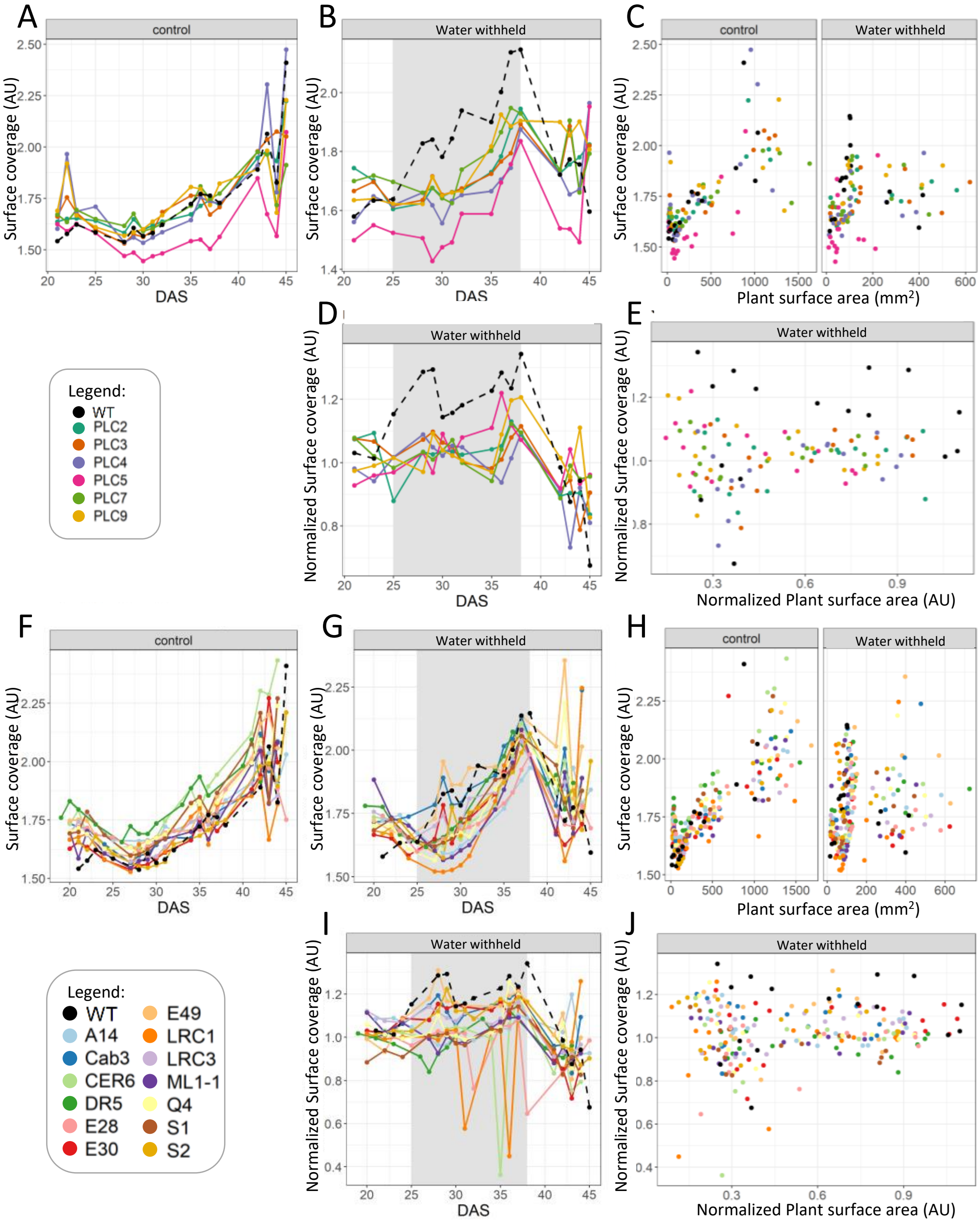


**Supplemental figure 25.** Surface coverage at control- and water limiting conditions.
The Surface coverage was measured for:
A, F) PLC-OE and TSEP grown at control conditions.
B, G) PLC-OE and TSEP at water-withholding conditions, with grey shading indicating period of water withholding (25-38 DAS).
C, H) PLC-OE and TSEP in Water-withholding conditions normalized to control per day
D, I ) PLC-OE and TSEP in Water withholding conditions normalized to control conditions.
E, J) PLC-OE and TSEP in water withholding conditions normalized to Projected plant surface area

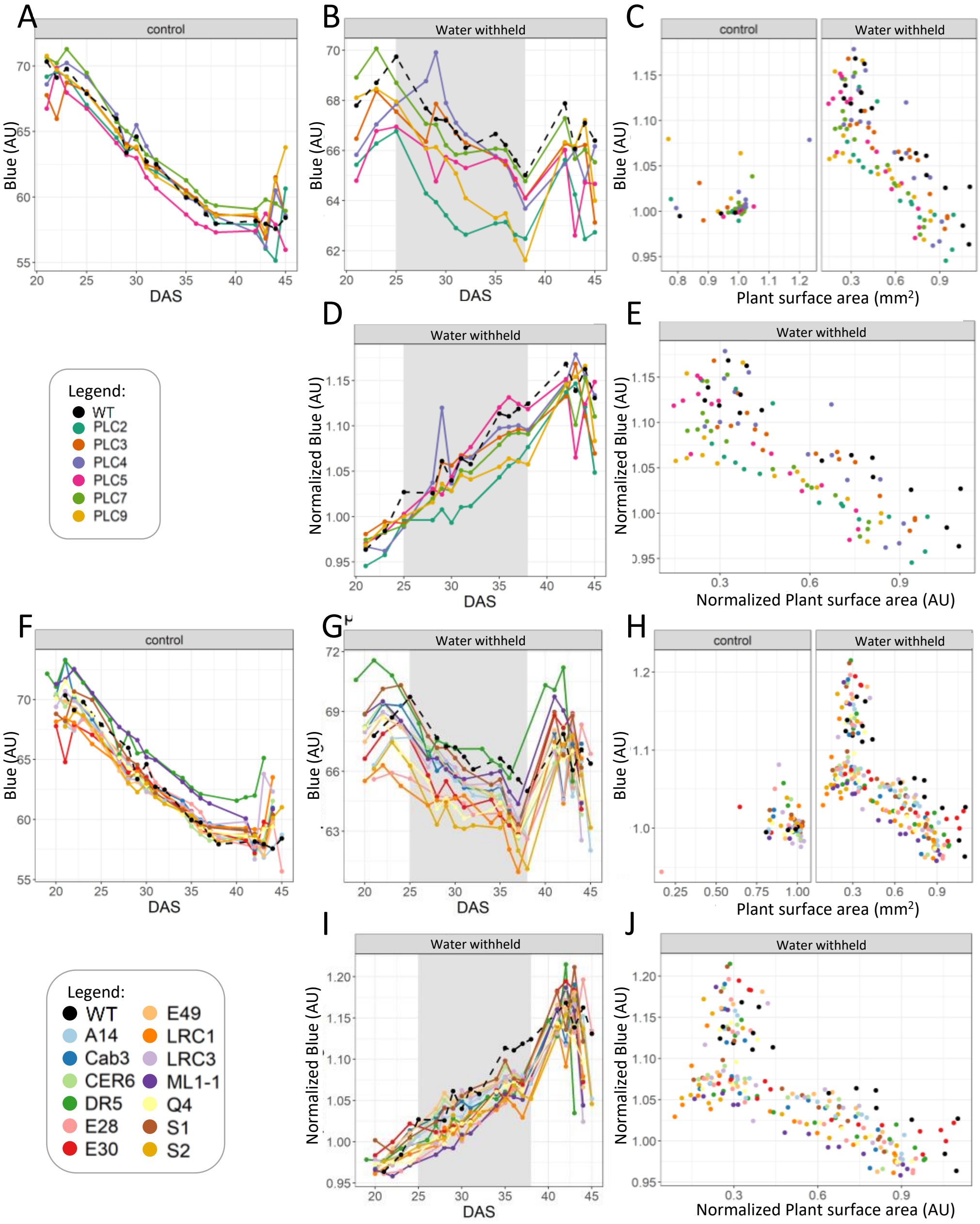


**Supplemental figure 26.** RGB Blue values at control- and water limiting conditions.
The RGB Blue values was measured for:
A, F) PLC-OE and TSEP grown at control conditions.
B, G) PLC-OE and TSEP at water-withholding conditions, with grey shading indicating period of water withholding (25-38 DAS).
C, H) PLC-OE and TSEP in Water-withholding conditions normalized to control per day
D, I ) PLC-OE and TSEP in Water withholding conditions normalized to control conditions.
E, J) PLC-OE and TSEP in water withholding conditions normalized to Projected plant surface area

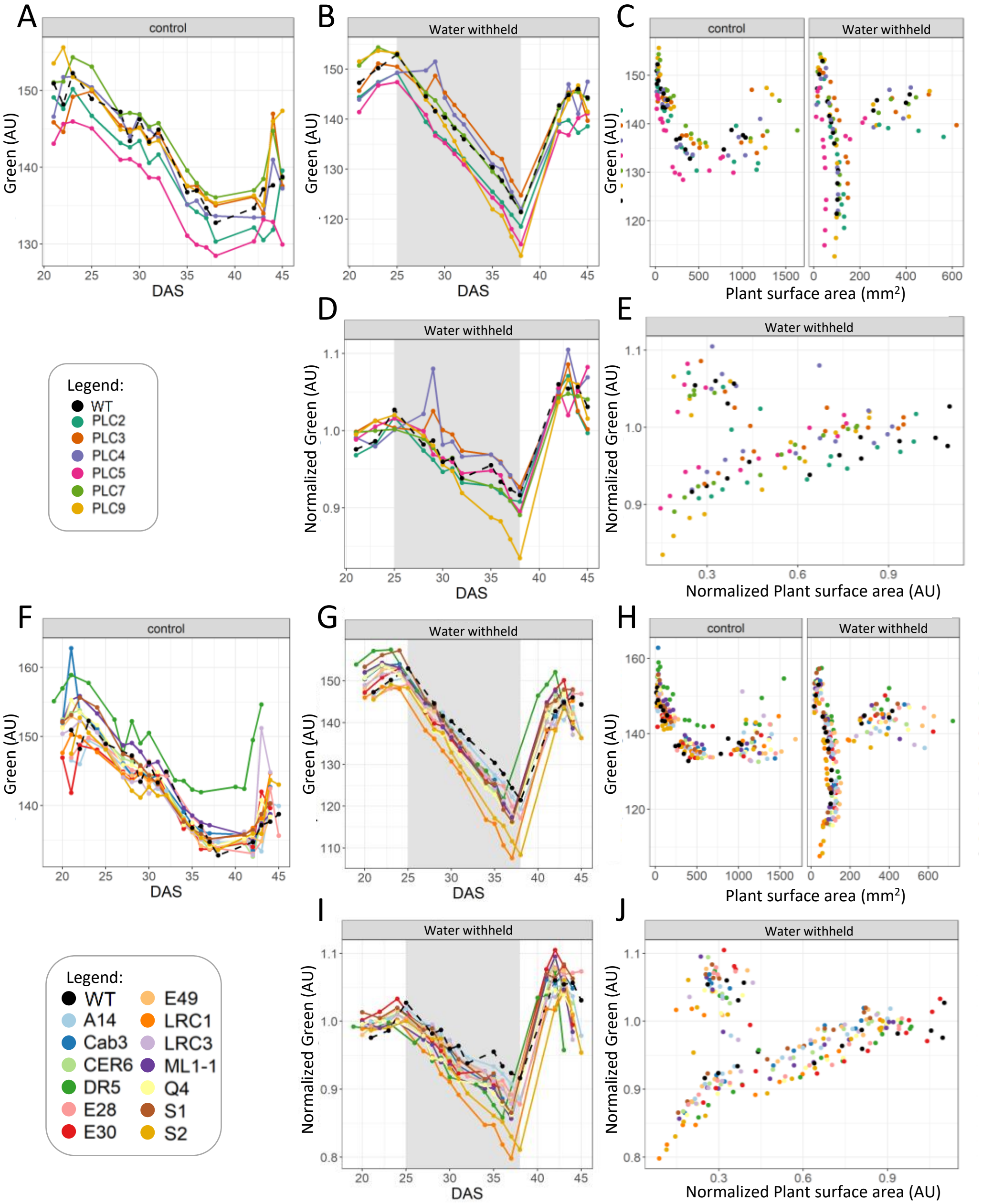


**Supplemental figure 27.** RGB Green values at control- and water limiting conditions. The RGB Green values was measured for:

A, F) PLC-OE and TSEP grown at control conditions.

B, G) PLC-OE and TSEP at water-withholding conditions, with grey shading indicating period of water withholding (25-38 DAS).

C, H) PLC-OE and TSEP in Water-withholding conditions normalized to control per day

D, I ) PLC-OE and TSEP in Water withholding conditions normalized to control conditions.

E, J) PLC-OE and TSEP in water withholding conditions normalized to Projected plant surface area

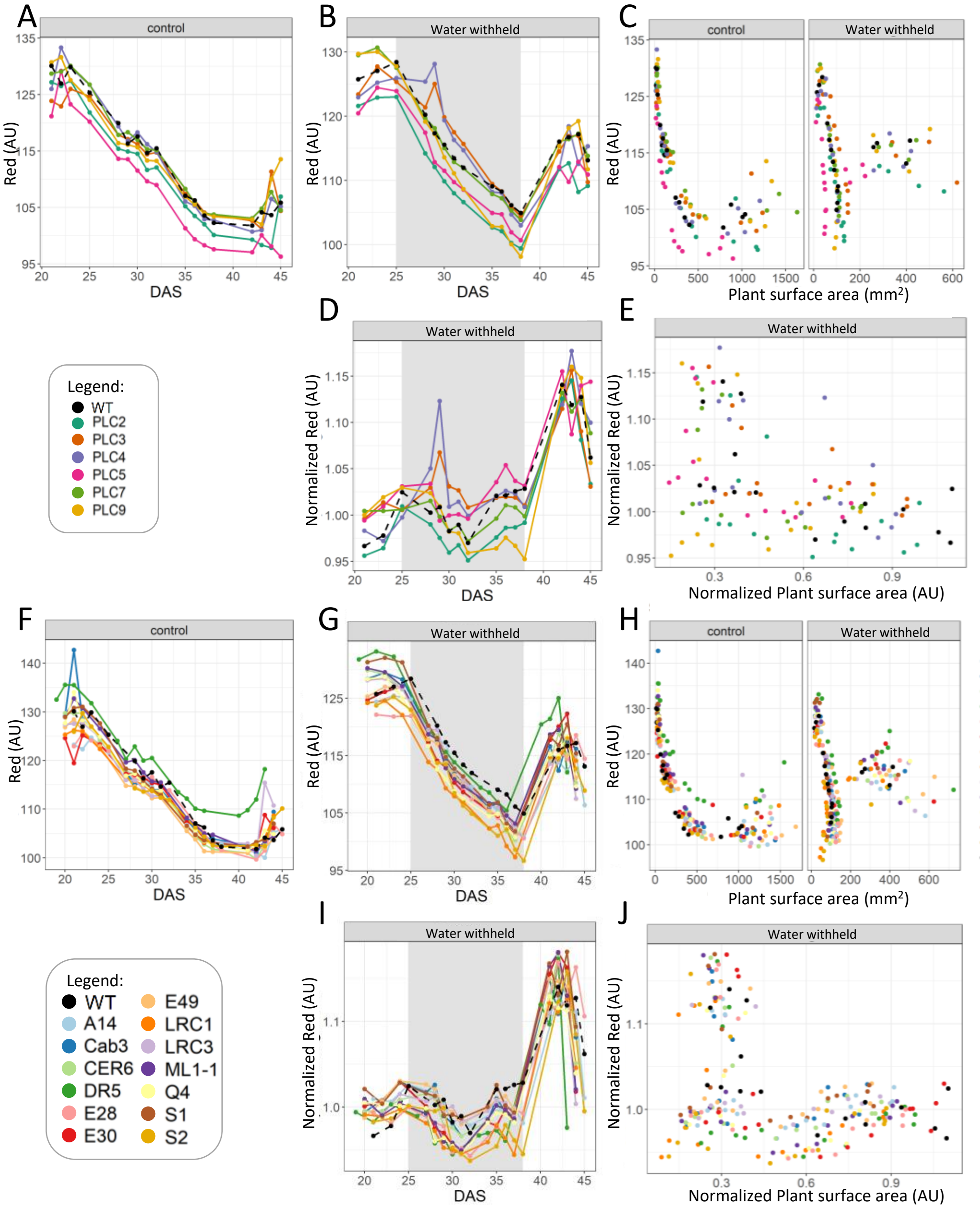


**Supplemental figure 28.** RGB Red values at control- and water limiting conditions.
The RGB Red values was measured for:
A, F) PLC-OE and TSEP grown at control conditions.
B, G) PLC-OE and TSEP at water-withholding conditions, with grey shading indicating period of water withholding (25-38 DAS).
C, H) PLC-OE and TSEP in Water-withholding conditions normalized to control per day
D, I ) PLC-OE and TSEP in Water withholding conditions normalized to control conditions.
E, J) PLC-OE and TSEP in water withholding conditions normalized to Projected plant surface area
